# Heightened Play following Play Observation in the Absence of 22KHz Calls in Juvenile Rats

**DOI:** 10.64898/2025.12.31.697143

**Authors:** Frédéric Michon Linde, Marijke Achterberg, Lena Valentina Kaufmann, Julian Packheiser, Sibi Bos, Kevin van Reenen, Valeria Gazzola, Christian Keysers

**Author notes:** Corresponding author: Frédéric Michon Linde.

## Abstract

Emotional contagion, the process leading to emotional state matching between individuals is considered a cornerstone for the proper functioning of social groups, via its contribution to group coordination and cohesion, the ability to learn from others and engage in prosocial behaviors. However, to date, most studies of emotional contagion investigate the transfer of negative emotional states thereby bypassing the neurobiology of positive affect sharing. In this study, we aimed to leverage on the innately rewarding and salient nature of play behaviour, to investigate the potential contagiousness of positive affective states in juvenile rats. Observers that had been moderately socially deprived for varying durations first witnessed highly-playful (Play) or non-playful (Control) demonstrator rats prior to being reunited and given the opportunity to freely interact. Surprisingly, we observed the emission of negatively-valenced 22 kHz calls in a subset of sessions which was also associated with heightened play behavior of the demonstrators. We found that the reunited observers showed an increase in play behavior following observation in the Play condition compared to the Control condition, but only after short social isolation and in sessions without 22 kHz calls. In addition, an overall higher number of positively-valenced 50 kHz calls were emitted in the Play condition, again in sessions without 22 kHz emissions. Despite the limitations of the experimental procedure, our results highlight the complex nature of positive emotional sharing and provide encouraging first indications of using social play observation to study positive emotional contagion.

**Highlights:**

- Social deprivation positively modulates play behavior and USV emission in reunited juvenile rats
- Heightened play is associated with 22 kHz USVs following social deprivation
- Heightened play was observed following play observation in the absence of 22 kHz USVs
- Play observation was associated with an overall increased number of 50 kHz calls emissions
- Play observation as an encouraging approach to study positive emotional contagion in rat

## Introduction

Emotional contagion, considered a basic form of empathy, leads to emotional state matching between individuals [1,2]. This process is thought to contribute to perceiving and evaluating the emotional state of others, coordinating behaviour, maintaining group cohesion, and engaging in prosocial behaviors [3,4]. Recent work in animal models has enabled us to deepen our understanding of the neuronal circuits underpinning this affective process [5–10]. However, most studies of emotional contagion investigate the transfer of negative emotional states, such as distress or pain [4,11,12], and there is a considerable lack of studies investigating the emotional contagion of positive emotions [13–16]. Developing rodent models of positive emotional sharing would be a valuable step towards characterizing the neural circuits involved in the contagion of another individual’s positive states and their potential differences with those involved in the contagion of negative emotional states [17].

One potential reason for the bias towards the study of negative affective states lies in the availability of behaviours (e.g. freezing and 22 kHz ultrasonic vocalizations [18–20]) that have been accepted as relatively reliable indicators of negative affective states. Reliable behaviors of positive emotional states, e.g.when obtaining a food reward, are less well established [21].

Social play also known as ‘play fighting’ or ‘rough-and-tumble play’ behaviour is an innately driven behaviour that is considered pleasurable [22], in a variety of species, including rats [23]. In juvenile rats, it is expressed vigorously [23–25]. Generally, a social play bout starts with one rat approaching the other rat and attempting to touch the nape of its neck, a behaviour known as pouncing. The solicited animal can respond in different ways, declining the invitation to play by walking away or rotating to its back with the other rat standing over it, resulting in the pin position. From this position, the solicited animal can launch a counterattack to the nape of the pinning rat which leads to an alternating interaction between the rats [22].

Most abundant between weaning and puberty [23–25], social play behavior is crucial for the development of social, emotional, and cognitive skills as well as the development of the brain [24–26]. Depriving rats of social play during the juvenile period (post-natal day 21-42) negatively impacts social and cognitive functions as well as brain physiology [27–30]. The rewarding properties of social play behaviour in rats have been demonstrated using place and operant conditioning tasks: rats spent more time in environments that have been associated with abundant social play [31,32] and they perform operant tasks for the opportunity to play [23,24]. The motivation to play can be modulated, as an increase in play behavior is observed in rats that have been isolated (up to 24 h) compared to rats that are socially housed [33–35]. In addition, playful interactions in rats are associated with 50 kHz ultrasonic vocalizations (USVs). These USVs are considered expressions of positive affect [36,37] as well as a means to coordinate social interactions, including play [38,39]. When exposed to playback of these 50 kHz USVs, rats respond by approaching the microphone [40–42], and are more inclined to press a lever for the opportunity to hear these USVs compared with 22 kHz calls [43]. Playback of 50 kHz calls further results in dopamine release [44] and increased activity in reward-related brain regions [45,46], suggesting that these calls can induce a positive affective state in conspecifics. Interestingly, play behavior can also be stimulated in rats by a direct social interaction with a more playful conspecific [47–49]. In these earlier paradigms, the focal rat could directly interact with a partner that varied in playfulness. Consequently, changes in the focal rat’s play behavior could reflect either *increased solicitation* by the more playful partner or *an enhanced motivation* to play arising from affective contagion.

To disentangle these components, we designed a setup in which direct physical interaction was prevented (see Figure 1). Two demonstrator rats that differed in their propensity to play were placed in one compartment, physically separated from two observer rats by a transparent, perforated divider allowing visual, auditory, and olfactory cues but no contact. Importantly, during the first 10 minutes of the observation phase, the two observer rats were also separated from each other by a second opaque divider. Thus, each observer could see the demonstrators but not its fellow observer. After this observation period, the divider between the observers was removed, allowing them to interact freely while still being separated from the demonstrators. This design allowed us to test whether observing demonstrators that engaged in higher versus lower levels of play would influence subsequent play behavior between the observers. We further manipulated the observers’ motivation for social interaction by varying the duration of their social isolation prior to test, as the effect of observation may depend on the observers being in a particular level of motivation to play. We predicted that both increased social deprivation and exposure to highly playful demonstrators would enhance the level of play displayed by the reunited observers, and thereby generate a novel paradigm to reveal emotional contagion of a positive affective state.

**Figure 1:**
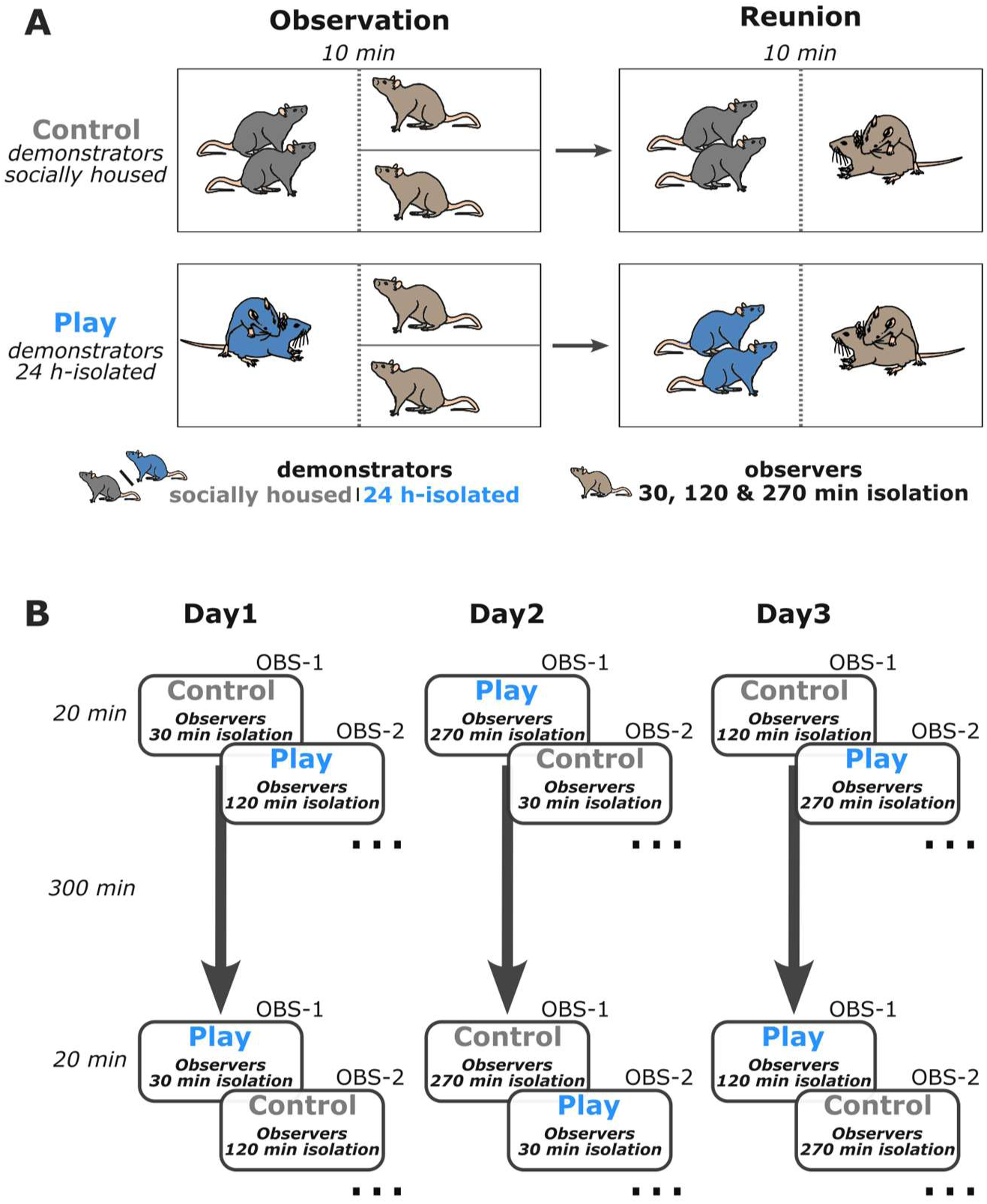
A within-subject paradigm to study the effect of play observation in juvenile rats. **A.** In a two phase paradigm, pairs of observer rats were first exposed to the social interactions, including social play, between two socially housed (Control observation) or 24 h-isolated demonstrators (Play observation) for 10 minutes, after which they were reunited. To incentivise different play-propensities of the observer animals across days, they were previously socially isolated for durations of 30, 120 or 270 min. **B.** Each of the 18 dyads of observers (OBS-1 .. OBS-18) were exposed to all conditions: 2 demonstrations (Play vs Control observation) x 3 social isolations (30 min, 120 min, 270 min) in a within-subject design within the course of 3 consecutive days. Within a single day, each observer dyad was exposed to both a Control and Play observation following the same isolation time with a delay of 300 min between experimental sessions. The order of presentation within a day (Play vs Control observation) and across the three days (30 min, 120 min, 270 min social isolation) was counterbalanced across all observer dyads.

## Materials and Methods

### 2.1 Subjects and housing

This study used juvenile male Wistar rats (n=84; Charles River, Sulzfeld, Germany). Upon arrival at 3 weeks of age, the animals were randomly allocated to a homecage (Macrolon type IV, 6 rats per cage with corn cob bedding under controlled conditions (22-24°C temperature, 55% relative humidity) at a reversed 12 h dark/light circle and with food and water available *ad libitum*. Rats were randomly assigned to one of three groups: Observers (n=36), Play demonstrators (n=36), and Control demonstrators (n=12). Each home cage housed 6 animals to which the same role was attributed (i.e. 6 Observers or 6 Play demonstrators or 6 Control demonstrators). Pairs of animals of the same homecage were further randomly assigned to a dyad which did not change throughout the experiment. In the experiment, the two observers or the two demonstrators were always cagemates, while observers and demonstrators always came from different cages. The main experiment was entirely within-subject, i.e. each observer dyad underwent all experimental conditions.

All experimental procedures were approved by the Centrale Commissie Dierproeven (CCD number: AVD8010020209724), the welfare body at the Netherlands Institute for Neuroscience (study dossier number: NIN181109) and in accordance with the Dutch and European regulations (Wet op Dierproeven 2014 and Guideline 86/609/EEC; Directive 2010/63/EU).

### 2.2 Apparatus

The experiment took place under red-lights, in a custom-made enclosure (l x w x h: 30.5 x 50.5 x 30 cm, Figure 1A) made of opaque black acrylic walls and of which the floor was covered with the same bedding material as in the home cages to enhance the animals’ comfort. The arena was divided into two equally-sized demonstrator and observer compartments by an acrylic divider, transparent and perforated, to let through olfactory, visual, and auditory cues. The observer compartment could be further divided in two with a removable opaque acrylic plate.

Behaviour was recorded with an overhead camera (acA1300-30gm, Basler) at 30 fps using Bonsai [50]. Three microphones (CM16/CMPA condenser ultrasound microphones, Avisoft) were hanged at the top of the walls in the observer and demonstrator compartments to record Ultrasonic vocalisations via an UltraSoundGate 116Hn audio recording system and the Avisoft-RECORDER software (Avisoft Bioacoustics, Glienicke, Germany).

### 2.3 Experimental procedure

Following arrival, the animals were first allowed to acclimate to their new environment for three days and were then handled 10 min in the stable for the three subsequent days. For three consecutive days prior to the experiment, all animals from each cage were habituated to their respective compartment for 20 min per day. Animals from different cages were never habituated simultaneously. The observers’ dyads were first habituated to be separated by the removable walls for 10 min, and reunited in the same compartment for the remaining 10min.

Rats underwent test sessions on three consecutive days in a within-subject design. Test sessions were divided in two phases, the observation phase and the reunion phase (Figure 1A). First, during the 10 min of observation phase, the observers’ dyads were separated in their compartment and witnessed the behaviour of either 24 h-isolated (Play observation) or socially housed (Control observation) demonstrators placed in the opposite compartment. The demonstrators were isolated for 24 h prior to the experiment or socially housed throughout the experiment in order to modulate the amount of play they displayed. Following observation, the opaque wall was removed to reunite the observer animals who were let to freely behave for a period of 10 min (reunion phase). The demonstrator animals remained in their compartment during the reunion phase. To modulate their motivation to play in the observers, observers had been isolated for 30,120 or 270 min prior to each session of the day.

Within the three days of testing, observers dyads thus went through all six combinations: Play or Control observations and prior isolation of 30 min, 120 min, or 270 min; in a pseudo-random, counterbalanced order (Figure 1B). On a single test day, one observer dyad experienced both Control and Play observation with a delay of 300 min and with the same prior isolation (i.e., if an observer dyad was isolated for 30 min before the first demonstrator exposure, they also received 30 min isolation before the second exposure). For each experimental session, the observers were exposed to a different demonstrator dyad to exclude a familiarization effect. Each play demonstrator dyad was socially isolated for 24 h and only used once per day, to ensure higher play levels. The control demonstrator dyads were tested three times each day, which is why three times more play demonstrators (n=36) were necessary than control demonstrators (n=12)

### 2.4 Analysis

#### 2.4.1 Behavioural characterization

Behavioural data were manually characterized using Observer XT16 and Ethovision XT9 software (Noldus Information Technology B.V., Wageningen, the Netherlands) by experimenters blinded to the experimental conditions. To characterize play behavior of both the observer and demonstrator dyads, pinning and pouncing - the two most characteristic behaviours in social play - were defined as previously described [22]:

- Pouncing: Snout or oral contact is directed to the nape of neck of the conspecific which is mostly followed by a rubbing movement
- Pinning: After contact with the nape, the animal holds the other animal down by using its front paws and standing over the animal. The other animal is lying on its back (supine position).

Play duration was quantified as the total amount of time spent pouncing or pinning for any animal of a dyad.

#### 2.4.2 Detections of ultrasonic vocalizations

50 kHz and 22 kHz ultrasonic vocalizations were separately detected using DeepSqueak [51] from a microphone located in the observers’ compartment. 50 kHz calls were detected using the *Rat Detector YoloR1* model, with 35 kHz and 70 kHz low and high frequency cutoff respectively. The 22 kHz emissions were detected with the *Long Rat Detector YOLO R1* model, and between 10 kHz to 30 kHz frequency cutoffs. The detections of the 22 kHz calls were further manually corrected by experimenters blind to the experimental conditions. Our recording setup did not enable us to identify the animal emitting the USVs, our analysis was thus limited to group effect at the scale of the sessions and between experimental conditions.

#### 2.4.3 Observer proximity analysis during observation

The observation compartment was divided in two equal zones (15 x 12 cm) to assess proximity of observers to the demonstrator compartment. This was quantified using EthoVision XT9 software (Noldus Information Technology, Wageningen, The Netherlands). The detection settings were center point detection and static subtraction with the animal being both brighter and darker than the background (bright contrast: 65-120, dark contrast: 25-85 pixel intensity units). Subject size was set between 355-30515 pixels. Smoothing was performed based on the 10 samples before and after every sample point. The time each observer rat spent in each zone was tracked over the 10 minutes observation phase and proximity was quantified as the percentage of time spent in the compartment half closest to the demonstrators.

#### 2.4.4 Observer anticipatory behaviour analysis during observation

Anticipatory behaviour of each observer rat during the 10 minutes observation phase was measured as described in [52] and expressed as behavioural switches per minute. The behaviors scored were: step, walk, run, jump, turn, explore, rear and groom. None of the individual behaviours differed between Control/Play sessions and social isolation time of the observers and because a general increase in activity is also considered a measure of anticipatory activity, it was decided to combine them and express them as behavioural switches per minute [53].

#### 2.4.5 Quantification and statistical analysis

Data were analysed using Python3 together with the scientific extension modules - numpy, pandas, statsmodels, matplotlib and seaborn - in addition to custom-written code, in Jupyter Notebooks. Bayesian models were computed in R using the brms package [54] and Bayes factors (BF_10_s) were computed using the bayestestR package [55].

Play duration of the demonstrators during the observation phase and of the observers during the reunion phase as well as the occurrence of ultrasonic vocalisations throughout the whole experimental sessions were quantified separately per experimental conditions (Control or Play) and for each observer isolation time (30, 120, 270 minutes) either at the level of the experimental phase (Observation and Reunion) or further broken down into minute long time bins.

All frequentist statistical analyses were performed using mixed linear models to test main and interaction fixed effects of the experimental conditions (Play and Control), observer isolation time (30, 120, or 270 minutes) and time within the sessions (minutes) on play duration, proximity occupancy, and ultrasonic calls occurrence, as below:

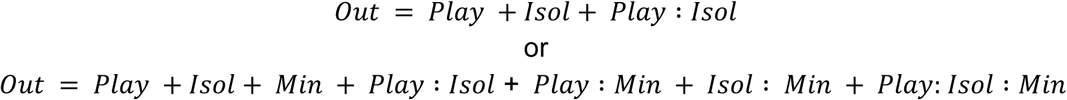

Where:

*Out*: outcome (Play duration, Pinning occurrence, Proximity, Anticipatory behavior or USVs occurrence)
*Play*: Play or Control experimental condition
*Isol*: Observers isolation time
*Min*: time bin, minute within phase

Experimental conditions (Control/Play observation and isolation duration) were considered as categorical factors and minutes throughout a phase as a continuous variable. To account for any potential variability in the propensity to play between each observer dyad, dyad identity was further used as a random effect (intercept). Statistical coefficient comparisons were then performed with a two-sided one-sample t-test against 0. Throughout the manuscripts, results from the mixed linear models are reported as: unstandardized coefficient(Β)[2.5%,97.5% confidence interval], t-test statistics, t-test p value.

For the Bayesian estimations, we used Cauchy priors centered around 0. The scaling of the priors were chosen based on the respective dependent variable (20s for play duration, 2 occurrences for pinning and behavioural switches, 10 occurrences for the 22kHz calls, and 50 for the 50kHz calls). BF_10_s of 1 indicate indifference for the support of either the null or alternative hypothesis. BF_10_s < 1 suggest evidence in favor of the null hypothesis whereas BF_10_s > 1 indicate evidence favoring the alternative hypothesis [56,57]. Using traditional bounds, we consider BF_10_<⅓ or BF_10_>3 as moderate evidence for the absence or presence of an effect, BF_10_<1/10 or BF_10_>10 as strong evidence, and 1/3<BF_10_<3 as merely circumstantial evidence.

## Results

### 3.1 Sessions in the Play condition are associated with ample play of the demonstrators and the emergence of 22 kHz call emissions

We first assessed whether demonstrators indeed showed differential amounts of play between the Play and Control observation. We quantified the total amount of time demonstrator animals spent playing during the observation phase (Figure 2A and Table 1). As expected, demonstrators spent a substantially longer amount of time playing in the Play observation condition compared with the control group (fixed effect of Play observation condition:Β_play_ 129s [92.3, 165.8], Z=6.88, p<0.001, BF_10_>100) and their play behavior did not systematically change over the 10 consecutive minutes of the observation period (fixed effect of minute:Β_minute_ 0.0s [−0.4, 0.5], Z=0.13, p=0.89, BF_10_=0.01; Supp. Figure 1 and Table 2).

**Figure 2:**
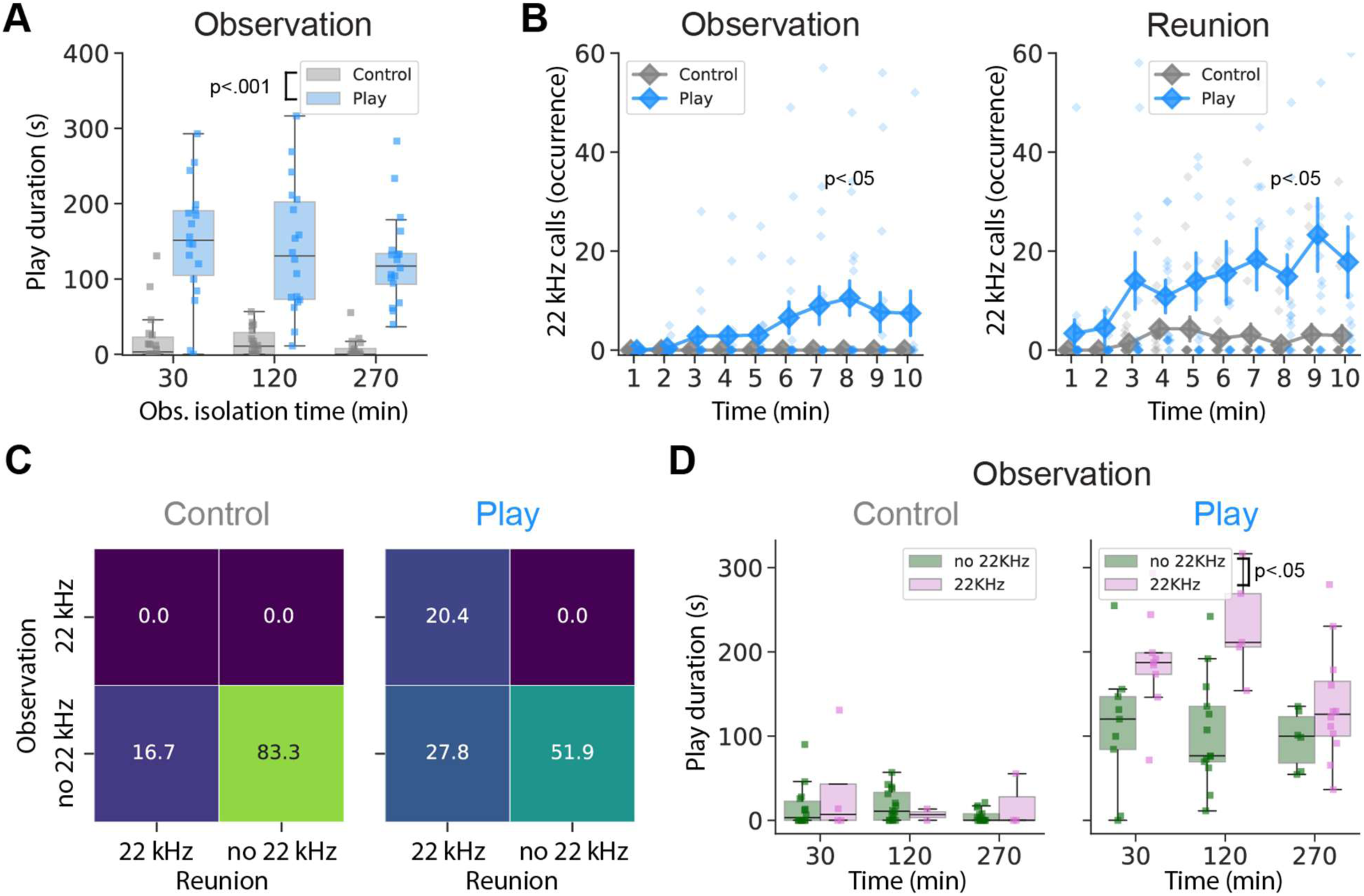
Emission of 22 kHz calls in sessions with longer play duration of demonstrators. **A.** Quantification of the total duration of social play behaviour displayed by the demonstrator dyads during the observation phase, separated by Control (socially housed demonstrators, grey bars) and Play observation (24 h-isolated demonstrators, blue bars) conditions and for the isolation time of the observer rats (30, 120 or 270 min prior to test). Boxplots represent median and quartiles, whiskers the rest of the data distribution outside of outliers and squares indicate values for single dyads. **B.** Quantification of the number of 22 kHz calls per minute, separated by Control (socially housed demonstrators, grey diamonds) and Play (24 h-isolated demonstrators, blue diamonds) conditions, during the observation phase (left panel) and the reunion phase (right panel). Connected diamonds represent the mean ± SEM, while non-connected diamonds indicate data from a single dyad. **C.** The distribution of sessions during which 22 kHz calls were detected, separated by experimental conditions and phases of the test. **D.** The total duration of play displayed by the demonstrators dyads during the observation phase, for sessions with 22 kHz calls detected (22 kHz, green bars) or not (no 22 kHz, pink bars), separated by Control (left panel) and Play (right panel) observation conditions and for isolation time of the observers (30, 120 and 270 min prior to test). Boxplots represent median and quartiles, whiskers the rest of the data distribution outside of outliers and squares indicate values for single dyads.

**1.** Demonstrators Play duration - Observation phase.

1- Demonstrators Play duration - Observation phase
|  | Coef. | Std.Err. | z | P>\|z\| | [0.025 | 0.975] | BF10 |
| --- | --- | --- | --- | --- | --- | --- | --- |
| Intercept | 20.325 | 13.385 | 1.519 | 0.129 | -5.908 | 46.559 | 0.72 |
| Isolation (120min) | -3.960 | 18.769 | -0.211 | 0.833 | -40.746 | 32.827 | 0.43 |
| Isolation(270min) | -13.068 | 18.769 | -0.696 | 0.486 | -49.855 | 23.718 | 0.68 |
| Play condition | 129.040 | 18.769 | 6.875 | 0.001 | 92.253 | 165.826 | > 100 |
| Isolation(120min) . Play condition | -5.686 | 26.543 | -0.214 | 0.830 | -57.710 | 46.338 | 0.50 |
| Isolation (270min) . Play condition | -13.211 | 26.543 | -0.498 | 0.619 | -65.235 | 38.813 | 0.63 |
| Observer dyads ID | 2.706 | 3.631 |  |  |  |  |  |

*Summary statistics of the mixed-linear model assessing the effect of observers isolation time, Play condition and their interaction on the amount of time demonstrators days spent playing in the observation phase.*

**2.** Demonstrators Play duration per mintues - Observation phase.

2- Demonstrators Play duration per mintues - Observation phase
|  | Coef. | Std.Err. | z | P>\|z\| | [0.025 | 0.975] | BF10 |
| --- | --- | --- | --- | --- | --- | --- | --- |
| Intercept | 2.051 | 2.321 | 0.884 | 0.377 | -2.498 | 6.601 | 0.01 |
| Isolation (120min) | -0.077 | 2.123 | -0.036 | 0.971 | -4.237 | 4.083 | 0.08 |
| Isolation(270min) | -1.336 | 2.123 | -0.629 | 0.529 | -5.497 | 2.824 | 0.11 |
| Play condition | 14.317 | 2.123 | 6.744 | 0.000 | 10.156 | 18.477 | > 100 |
| Isolation(120min) . Play condition | -3.414 | 3.002 | -1.137 | 0.255 | -9.298 | 2.470 | 0.21 |
| Isolation (270min) . Play condition | -1.339 | 3.002 | -0.446 | 0.656 | -7.223 | 4.545 | 0.11 |
| minute | 0.032 | 0.242 | 0.133 | 0.894 | -0.442 | 0.506 | 0.01 |
| Isolation (120min) . minute | -0.058 | 0.342 | -0.170 | 0.865 | -0.729 | 0.613 | 0.02 |
| Isolation(270min) . minute | 0.005 | 0.342 | 0.016 | 0.988 | -0.665 | 0.676 | 0.01 |
| Minute . Play condition | -0.257 | 0.342 | -0.751 | 0.453 | -0.927 | 0.414 | 0.02 |
| Isolation(120min) . minute . Play condition | 0.517 | 0.484 | 1.069 | 0.285 | -0.431 | 1.466 | 0.03 |
| Isolation (270min) . minute . Play condition | 0.003 | 0.484 | 0.007 | 0.995 | -0.945 | 0.951 | 0.02 |
| Observer dyads ID | 2.989 | 0.143 |  |  |  |  |  |

*Summary statistics of the mixed-linear model assessing the effect of observers isolation time, Play condition, minute within the phase and their interaction on the amount of time demonstrators days spent playing in the observation phase.*

The amount of play demonstrators engaged in was not influenced by the observers’ state as we found moderate to strong evidence of absence of an effect of the observers’ isolation time (increase relative to 30 min isolation time: −4s [−40.8, 32.8], Z=−0.21, p=0.83, BF_10_=0.08 and - 13.1s [−49.9, 23.8], Z=−0.70, p=0.49, BF_10_=0.11; for 120 min and 270 min isolation time respectively).

While the modulation of the demonstrators’ play behavior followed the expected outcome, when we turned to the ultrasonic vocalizations emitted by the rats, we noticed that some sessions were associated with the emission of 22 kHz calls (Figure 2B and Suppl. Figure 3). These emerged prominently in association with the Play observation condition during the observation phase, for which we found a significant interaction between time throughout the session and Play observation condition (+0.60 occurrences [0.25, 0.97] every minute, Z=3.30, p<0.001, BF_10_=3.56; Figure 3A and Table 3). The increased occurrence of 22 kHz calls over time did not substantially differ between the observers’ isolation times (Β_120min.minute_:-0.19 occurrences [−0.727,0.467], Z=−0.70, p=0.32, BF_10_=0.03; Β_270min.minute_:-0.57 occurrences [−1.08, −0.06], Z=−2.18, p=0.03, BF_10_=0.22). We observed a similar phenomenon during the reunion phase (+0.68 occurrences [0.07, 1.28] every minute, Z=2.20, p<0.02, BF_10_=0.40) without difference across observers’ isolation time (Figure 2B and Supp Figure 3, Table 3-4).

**Figure 3:**
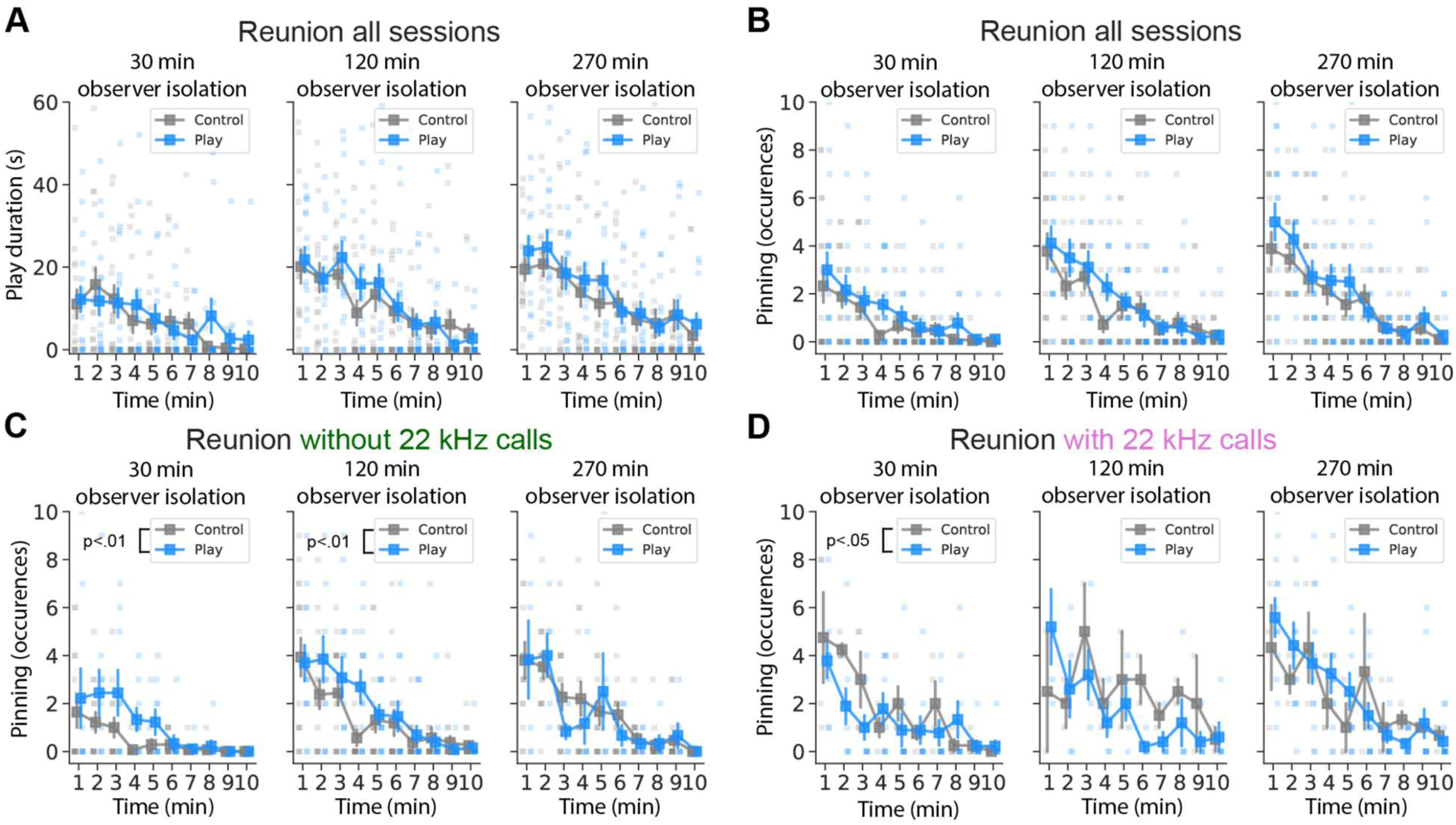
Enhanced play behavior following Play observation, only with mild social deprivation and in the absence of 22 kHz calls. **A.** Quantification of the total time spent playing by the observer dyads per minute in the reunion phase, following observation of Control (socially housed demonstrators, grey squares) and Play (24 h-isolated demonstrators, blue squares) observation and per isolation time of the observers (30, 120 and 270 minutes prior to test) **B-C-D.** Quantification of the number of pins per minute by observer dyads during the reunion phase following observation of Control (socially housed demonstrators, grey squares) and Play (24 h-isolated demonstrators, blue squares) observation and per isolation time of the observers (30, 120 and 270 min prior to test) (B), in sessions without 22 kHz calls detected (C) and in session with 22 kHz calls detected (D). For all panels, connected squares represent the mean ± SEM. Non-connected squares represent data from individual dyads.

**3.** 22KHz calls - Observation phase.

3- 22KHz calls - Observation phase
|  | Coef. | Std.Err. | z | P>\|z\| | [0.025 | 0.975] | BF10 |
| --- | --- | --- | --- | --- | --- | --- | --- |
| Intercept | -0.172 | 1.642 | -0.105 | 0.917 | -3.391 | 3.047 | 0.00 |
| Isolation (120min) | 0.000 | 1.146 | 0.000 | 1.000 | -2.246 | 2.246 | 0.09 |
| Isolation(270min) | 0.000 | 1.146 | 0.000 | 1.000 | -2.246 | 2.246 | 0.08 |
| Play condition | -0.959 | 1.146 | -0.837 | 0.403 | -3.205 | 1.287 | 0.11 |
| Isolation(120min) . Play condition | 0.478 | 1.621 | 0.295 | 0.768 | -2.699 | 3.654 | 0.13 |
| Isolation (270min) . Play condition | 1.515 | 1.621 | 0.935 | 0.350 | -1.662 | 4.691 | 0.18 |
| minute | 0.000 | 0.131 | 0.000 | 1.000 | -0.256 | 0.256 | 0.01 |
| Isolation (120min) . minute | -0.000 | 0.185 | -0.000 | 1.000 | -0.362 | 0.362 | 0.01 |
| Isolation(270min) . minute | -0.000 | 0.185 | -0.000 | 1.000 | -0.362 | 0.362 | 0.01 |
| Minute . Play condition | 0.612 | 0.185 | 3.312 | 0.001 | 0.250 | 0.974 | 3.56 |
| Isolation(120min) . minute . Play condition | -0.190 | 0.261 | -0.727 | 0.467 | -0.702 | 0.322 | 0.03 |
| Isolation (270min) . minute . Play condition | -0.568 | 0.261 | -2.176 | 0.030 | -1.080 | -0.056 | 0.22 |
| Observer dyads ID | 1.956 | 0.169 |  |  |  |  |  |

Summary statistics of the mixed-linear model assessing the effect of observers isolation time, Play condition, minute within the phase and their interaction on the number of 22 kHz calls emissions in the observation phase.

**4.** 22KHz calls - Reunion phase.

4- 22KHz calls - Reunion phase
|  | Coef. | Std.Err. | z | P>\|z\| | [0.025 | 0.975] | BF10 |
| --- | --- | --- | --- | --- | --- | --- | --- |
| Intercept | 1.259 | 4.786 | 0.263 | 0.792 | -8.121 | 10.639 | 0.00 |
| Isolation (120min) | -0.816 | 4.880 | -0.167 | 0.867 | -10.380 | 8.748 | 0.30 |
| Isolation(270min) | -4.177 | 4.880 | -0.856 | 0.392 | -13.741 | 5.387 | 0.42 |
| Play condition | -5.589 | 4.880 | -1.145 | 0.252 | -15.153 | 3.975 | 0.51 |
| Isolation(120min) . Play condition | 3.911 | 6.901 | 0.567 | 0.571 | -9.614 | 17.437 | 0.36 |
| Isolation (270min) . Play condition | 0.999 | 6.901 | 0.145 | 0.885 | -12.526 | 14.524 | 0.35 |
| minute | 0.005 | 0.219 | 0.022 | 0.983 | -0.424 | 0.434 | 0.01 |
| Isolation (120min) . minute | -0.016 | 0.310 | -0.051 | 0.959 | -0.623 | 0.591 | 0.02 |
| Isolation(270min) . minute | 0.256 | 0.310 | 0.828 | 0.408 | -0.350 | 0.863 | 0.02 |
| Minute . Play condition | 0.678 | 0.310 | 2.190 | 0.029 | 0.071 | 1.284 | 0.40 |
| Isolation(120min) . minute . Play condition | -0.389 | 0.438 | -0.888 | 0.374 | -1.247 | 0.469 | 0.03 |
| Isolation (270min) . minute . Play condition | -0.145 | 0.438 | -0.332 | 0.740 | -1.003 | 0.713 | 0.02 |
| Observer dyads ID | 10.626 | 0.491 |  |  |  |  |  |

Summary statistics of the mixed-linear model assessing the effect of observers isolation time, Play condition, minute within the phase and their interaction on the number of 22 kHz calls emissions in the reunion phase.

When quantifying sessions during which at least one 22 kHz call was detected, we observed that 22 kHz calls were likely associated with the reunion of rats, demonstrators and observers alike, who had been socially isolated, and thus likely emitted these calls during play behavior, either while engaging in or observing it (Figure 2C). We found that during the observation phase, none were detected in the Control condition and in 20.4% (n=11) of the Play observation sessions. A larger percentage of sessions with 22 kHz calls were found during the reunion phase, respectively 16.4% (n=9) and 27.8% (n=15) in the Control and Play observation conditions. Only 16.0% (n=3) of the observer dyads were never associated to a session with 22 kHz calls emitted while it was the case for 55.0% (n=10) of the 24 h isolated demonstrators’ dyads, suggesting that 22 kHz calls emissions were more tightly associated to demonstrators’ than observers’ identity.

Because 22 kHz calls are considered the expression of negative affect in rats [18–20,36] and play is suppressed by negative factors such as food deprivation and intense light conditions during testing [24,52,58,59], we reasoned that play behaviour could be influenced in sessions associated with the emission of negative calls. To test this hypothesis, we separated sessions based on the occurrence of 22 kHz calls at any phase (22 kHz sessions or no 22 kHz sessions) and quantified the amount of time the demonstrator dyads spent playing during the observation phase (Figure 2D). Although we did not find any difference in play duration in the Control observation condition (Β_22kHz presence_: +19.70s [−46.90, 7.50], Z=1.42, p=0.16, BF_10_=0.58, Table 4), demonstrator dyads played longer in the Play observation condition with 22 kHz emissions than in those without (Β_22kHz presence_: +76.90s [−136.90, −16.90], Z=2.51, p=0.01, BF_10_=18.93; Table 5-6). Thus sessions with longer play duration of the demonstrators were associated with emissions of 22kHz calls (Figure 2D).

**5.**
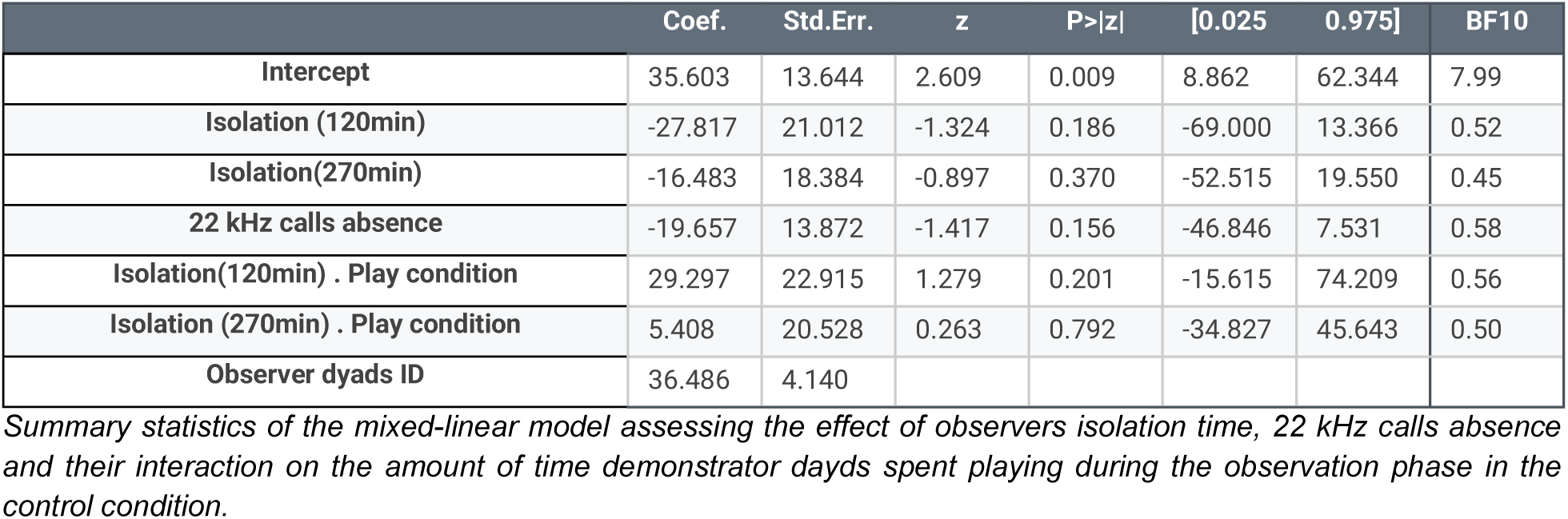
Demonstrators Play duration - 22KHz split - Control.

**6.** Demonstrators Play duration - 22KHz split - Play.

6- Demonstrators Play duration - 22KHz split - Play
|  | Coef. | Std.Err. | z | P>\|z\| | [0.025 | 0.975] | BF10 |
| --- | --- | --- | --- | --- | --- | --- | --- |
| Intercept | 187.814 | 21.644 | 8.677 | 0.000 | 145.392 | 230.236 | > 100 |
| Isolation (120min) | 43.471 | 36.183 | 1.201 | 0.230 | -27.447 | 114.390 | 0.80 |
| Isolation(270min) | -51.306 | 28.674 | -1.789 | 0.074 | -107.507 | 4.895 | 1.65 |
| 22 kHz calls absence | -76.909 | 30.595 | -2.514 | 0.012 | -136.874 | -16.944 | 18.93 |
| Isolation(120min) . Play condition | -49.882 | 45.832 | -1.088 | 0.276 | -139.712 | 39.948 | 1.04 |
| Isolation (270min) . Play condition | 36.626 | 45.040 | 0.813 | 0.416 | -51.651 | 124.904 | 0.76 |
| Observer dyads ID | 0.000 | 11.057 |  |  |  |  |  |

Summary statistics of the mixed-linear model assessing the effect of observers isolation time, 22 kHz calls absence and their interaction on the amount of time demonstrator days spent playing during the observation phase in the play condition.

We then assessed whether the demonstrators’ behaviour influenced that of the observers during the observation phases. Observer rats spent the majority of their time in the half of their observer compartment closest to the demonstrators (one-sample T-test against 50%, sessions without 22 kHz calls: mean=76.01% [73.3, 78.7] 95% confidence interval, T=19.15, p<0.001, BF_10_>100; sessions with 22 kHz calls: mean=75.81% [71.35, 80.28] 95% confidence interval, T=11.53, p<0.001, BF_10_>100). However, we found evidence for the absence of an effect of Play vs Control observation both for sessions with and without 22 kHz detected (+5.68% [−4.90, 16.20], Z=1.05, p=0.29, BF_10_=0.21 and +0.01% [−12.30, 12.50], Z=0.02, p=0.99, BF_10_=0.25, respectively; Supp Figure 2, Table 15-16). In addition, anticipatory behaviors did not differ betweenPlay vs Control observations, with evidence for the absence of an effect (−0.08 switches/min [−1.55, 1.39], Z=−0.11, p=0.91, BF_10_=0.12 and −0.38 switches/min [−2.08, 1.32], Z=−0.44, p=0.66, BF_10_=0.17 for sessions with and without 22 kHz calls emitted, respectively; Supp Figure 2, Table 17-18). This shows that social observation is salient and attractive, independently of whether the demonstrators are involved in play.

### 3.2 Increased play behavior following short social isolation and in the absence of 22 kHz call emissions

Next, to investigate whether observer rats were emotionally influenced by the behaviours of the demonstrators, we turned to the reunion phase and the behaviors of the observer dyads. In our paradigm we predicted an increase of play behavior due to the duration of the social isolation of the observers and, if playful state contagion occurred, an additional increase following play observation compared to the control observation.

We first quantified the duration engaged in play per minute of the reunion phase (Figure 3A and Table 7). As predicted, observer rats engaged in more play following 120 and 270 min of social isolation compared to 30 min (respectively, +5.50s [0.40, 10.50] and +7s [1.90, 12.00] per minute; Z=2.10, p=0.04, BF_10_=1.22, and Z=2.68, p=0.007, BF_10_=5.39). We also found a significant effect of time within the sessions. For all isolation times, the observers decreased the amount of time playing over the session (−1.60s [−2.20, −1.00] per minute Z=−5.43, p<0.0001, BF_10_=76.76). However we found evidence of absence of an effect of observing Play vs Control demonstrators (Β_Play observation_: −1.60s [−6.70, 3.50] per minute, Z=−0.60, p=0.54, BF_10_=0.03).

**7.** Observers Play duration - Reunion Phase.

7- Observers Play duration - Reunion Phase
|  | Coef. | Std.Err. | z | P>\|z\| | [0.025 | 0.975] | BF10 |
| --- | --- | --- | --- | --- | --- | --- | --- |
| Intercept | 22.650 | 9.808 | 2.309 | 0.021 | 3.427 | 41.874 | > 100 |
| Isolation (120min) | 5.451 | 2.602 | 2.095 | 0.036 | 0.352 | 10.550 | 1.22 |
| Isolation(270min) | 6.971 | 2.602 | 2.680 | 0.007 | 1.872 | 12.070 | 5.39 |
| Play condition | -1.587 | 2.602 | -0.610 | 0.542 | -6.686 | 3.512 | 0.12 |
| Isolation(120min) . Play condition | 5.860 | 3.679 | 1.593 | 0.111 | -1.351 | 13.071 | 0.46 |
| Isolation (270min) . Play condition | 5.372 | 3.679 | 1.460 | 0.144 | -1.839 | 12.583 | 0.43 |
| minute | -1.609 | 0.296 | -5.426 | 0.000 | -2.190 | -1.027 | 76.76 |
| Isolation (120min) . minute | -0.219 | 0.419 | -0.523 | 0.601 | -1.041 | 0.602 | 0.02 |
| Isolation(270min) . minute | -0.292 | 0.419 | -0.696 | 0.486 | -1.114 | 0.530 | 0.02 |
| Minute . Play condition | 0.418 | 0.419 | 0.998 | 0.318 | -0.404 | 1.240 | 0.03 |
| Isolation(120min) . minute . Play condition | -0.998 | 0.593 | -1.683 | 0.092 | -2.160 | 0.164 | 0.10 |
| Isolation (270min) . minute . Play condition | -0.774 | 0.593 | -1.306 | 0.192 | -1.937 | 0.388 | 0.04 |
| Observer dyads ID | 90.455 | 3.549 |  |  |  |  |  |

Summary statistics of the mixed-linear model assessing the effect of observers isolation time, Play condition, minute within the phase and their interaction on the amount of time observer days spent playing in the reunion phase.

We next reasoned that play contagion might increase willingness of the two members of a dyad to engage in play together, in which case an effect of play observation could be more visible when looking at the specific occurrence of pins (Figure 3B and Table 8). As for play duration, we did observe an increased number of pins per minute following 120 and 270 min of social isolation (respectively, +1.20 occurrences [0.35, 2.00] and +1.90 occurrences [1.10, 2.80] per minute; Z=2.80, p=0.005, BF_10_=8.22 and Z=4.63, p<0.001, BF_10_>100). In addition, a decrease in the number of pins over minutes within the phase was found (−0.24 occurrences [−0.34, −0.15] per minute Z=−5.11, p<0.001, BF_10_>100). However, we did not observe an effect of Play observation on the number of pins (+0.68 occurrences [−0.14, 1.50] per minute, Z=1.63, p=0.10, BF_10_=0.58).

**8.**
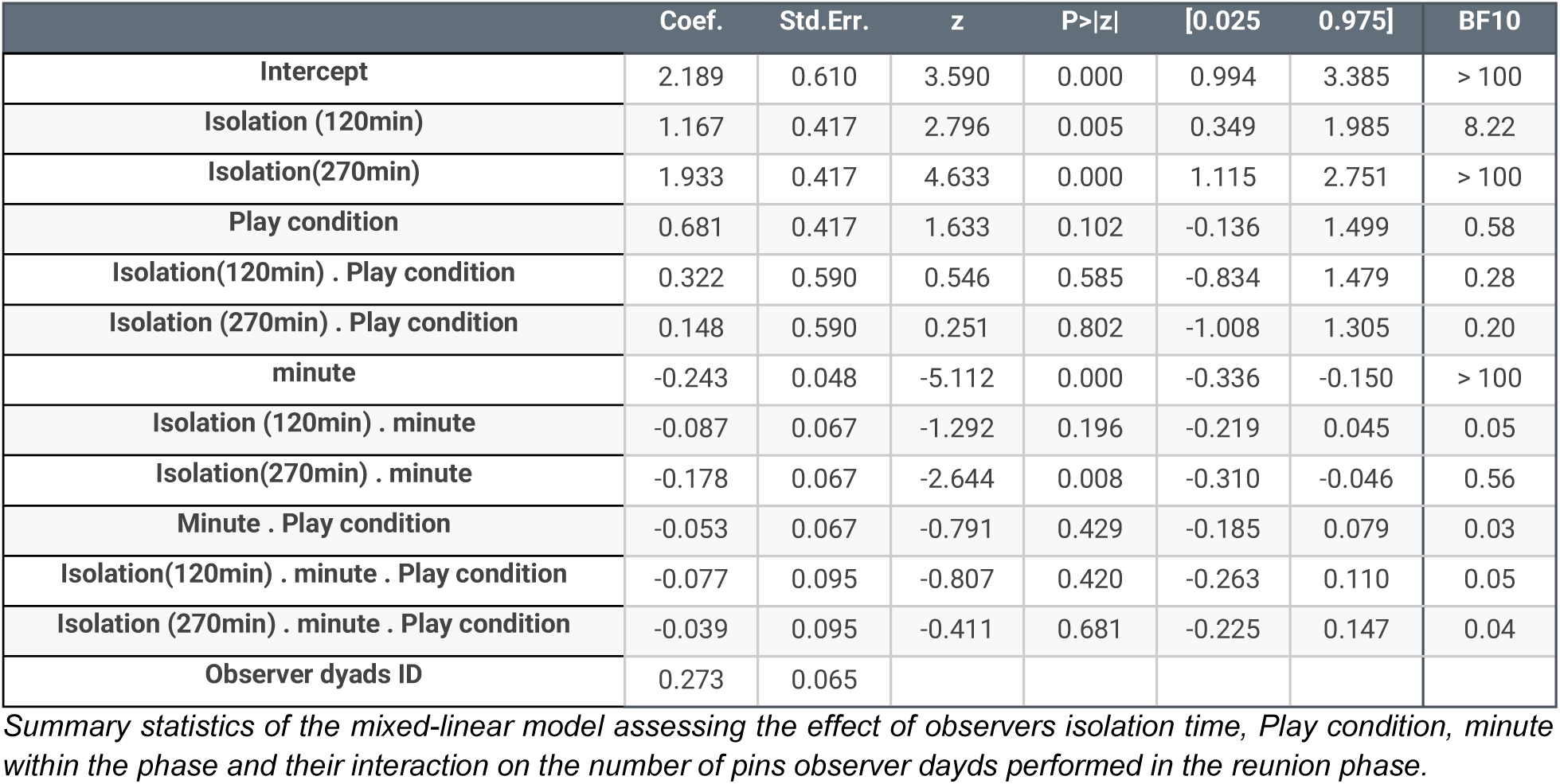
Observers Pinning occurrence - Reunion Phase.

8- Observers Pinning occurrence - Reunion Phase
|  | Coef. | Std.Err. | z | P>\|z\| | [0.025 | 0.975] | BF10 |
| --- | --- | --- | --- | --- | --- | --- | --- |
| Intercept | 2.189 | 0.610 | 3.590 | 0.000 | 0.994 | 3.385 | > 100 |
| Isolation (120min) | 1.167 | 0.417 | 2.796 | 0.005 | 0.349 | 1.985 | 8.22 |
| Isolation(270min) | 1.933 | 0.417 | 4.633 | 0.000 | 1.115 | 2.751 | > 100 |
| Play condition | 0.681 | 0.417 | 1.633 | 0.102 | -0.136 | 1.499 | 0.58 |
| Isolation(120min) . Play condition | 0.322 | 0.590 | 0.546 | 0.585 | -0.834 | 1.479 | 0.28 |
| Isolation (270min) . Play condition | 0.148 | 0.590 | 0.251 | 0.802 | -1.008 | 1.305 | 0.20 |
| minute | -0.243 | 0.048 | -5.112 | 0.000 | -0.336 | -0.150 | > 100 |
| Isolation (120min) . minute | -0.087 | 0.067 | -1.292 | 0.196 | -0.219 | 0.045 | 0.05 |
| Isolation(270min) . minute | -0.178 | 0.067 | -2.644 | 0.008 | -0.310 | -0.046 | 0.56 |
| Minute . Play condition | -0.053 | 0.067 | -0.791 | 0.429 | -0.185 | 0.079 | 0.03 |
| Isolation(120min) . minute . Play condition | -0.077 | 0.095 | -0.807 | 0.420 | -0.263 | 0.110 | 0.05 |
| Isolation (270min) . minute . Play condition | -0.039 | 0.095 | -0.411 | 0.681 | -0.225 | 0.147 | 0.04 |
| Observer dyads ID | 0.273 | 0.065 |  |  |  |  |  |

Summary statistics of the mixed-linear model assessing the effect of observers isolation time, Play condition, minute within the phase and their interaction on the number of pins observer days performed in the reunion phase.

Given the association of 22 kHz and play behavior in demonstrators, we then assessed observer play behavior separately for sessions without and with 22 kHz calls detected (Figure 3C-D). For sessions without any 22 kHz call detected, we again found a similar effect of the observers’ isolation time and time spent during the phase on pin occurrence (Figure 3C and Table 9). Importantly, for these sessions Play observation was followed by an increased number of pins (+1.4 occurrences [0.41,2.40] per minute, Z=2.75, p=0.006, BF_10_=3.56). We found similar trends when quantifying the total amount of time spent playing (Supp Figure 4A and Table 10), suggesting that the number of pins is a more sensitive read out of the effect of Play observation on play behavior. We then tested whether the total number of pins displayed by observers following Play observation, was directly related to the amount of play that had witnessed being displayed by the demonstrator animals during the observation phase, but found evidence for the absence of such an effect (Β_Play demonstrators_: 0.035 occurence [−0.08, 0.15], Z=0.59, p=0.55, BF_10_=0.03).

**Figure 4:**
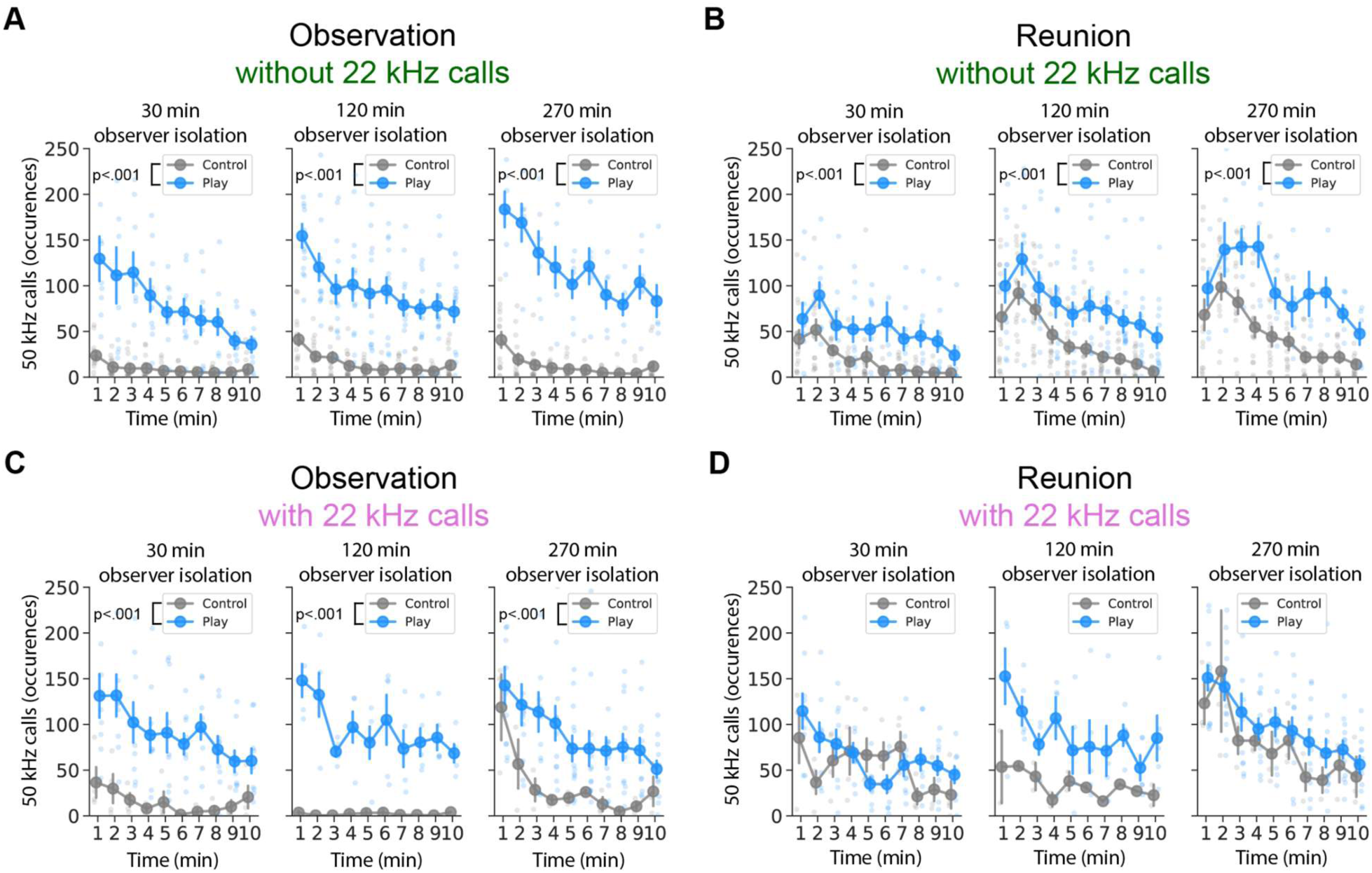
Enhanced 50 kHz calls emission associated with Play observation. Quantification of the number of 50 kHz calls emissions per minute, separated by the control and play condition and isolation time of the observers (30, 120 and 270 min prior to test): in the absence of 22 kHz calls during the observation (A) and the reunion phase (B); as well as in the presence of 22 kHz calls during the observation (C) and reunion phase (D). For all panels, connected circles represent the mean ± SEM. Non-connected circles represent data from individual dyads.

**9.**
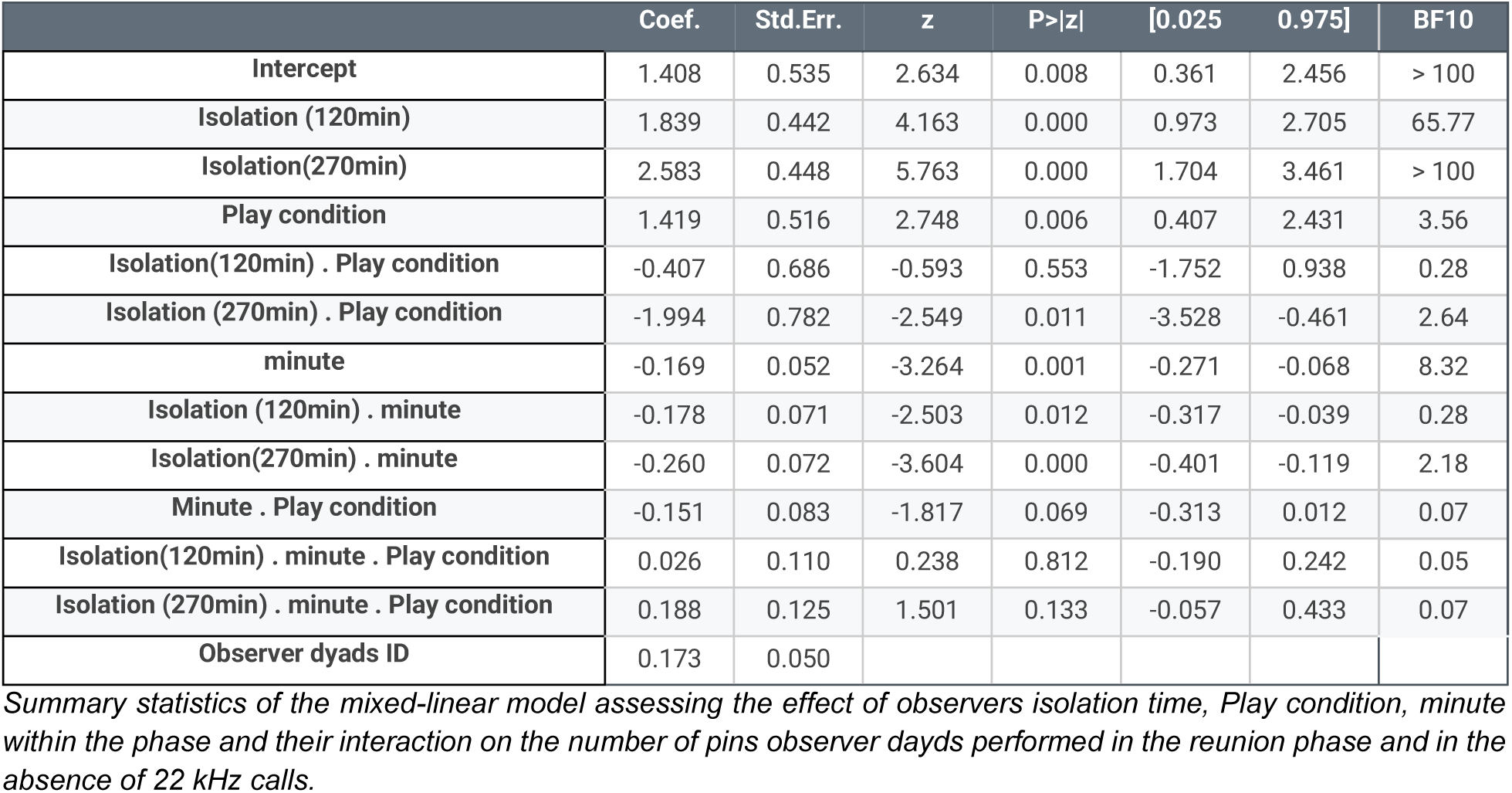
Observers Pinning occurrence - Reunion Phase - no 22KHz.

**10.** Observers Pinning occurrence - Reunion Phase - 22KHz.

10- Observers Pinning occurrence - Reunion Phase - 22KHz
|  | Coef. | Std.Err. | z | P>\|z\| | [0.025 | 0.975] | BF10 |
| --- | --- | --- | --- | --- | --- | --- | --- |
| Intercept | 4.575 | 1.021 | 4.480 | 0.000 | 2.574 | 6.577 | > 100 |
| Isolation (120min) | -1.168 | 1.125 | -1.038 | 0.299 | -3.374 | 1.037 | 0.38 |
| Isolation(270min) | -0.148 | 0.989 | -0.149 | 0.881 | -2.086 | 1.791 | 0.39 |
| Play condition | -1.752 | 0.785 | -2.232 | 0.026 | -3.290 | -0.213 | 0.87 |
| Isolation(120min) . Play condition | 2.735 | 1.345 | 2.033 | 0.042 | 0.098 | 5.372 | 1.01 |
| Isolation (270min) . Play condition | 2.950 | 1.146 | 2.574 | 0.010 | 0.704 | 5.196 | 3.34 |
| minute | -0.502 | 0.103 | -4.847 | 0.000 | -0.704 | -0.299 | > 100 |
| Isolation (120min) . minute | 0.308 | 0.179 | 1.716 | 0.086 | -0.044 | 0.659 | 0.11 |
| Isolation(270min) . minute | 0.122 | 0.158 | 0.770 | 0.441 | -0.188 | 0.431 | 0.05 |
| Minute . Play condition | 0.229 | 0.124 | 1.840 | 0.066 | -0.015 | 0.473 | 0.39 |
| Isolation(120min) . minute . Play condition | -0.465 | 0.213 | -2.182 | 0.029 | -0.883 | -0.047 | 0.22 |
| Isolation (270min) . minute . Play condition | -0.423 | 0.183 | -2.316 | 0.021 | -0.780 | -0.065 | 0.28 |
| Observer dyads ID | 0.580 | 0.176 |  |  |  |  |  |

Summary statistics of the mixed-linear model assessing the effect of observers isolation time, Play condition, minute within the phase and their interaction on the number of pins observer dayds performed playing in the reunion phase and in the presence of 22 kHz calls.

In sessions with 22 kHz calls, we found no evidence for an effect of Play observation (−1.75 occurrences [−3.29, −0.21] per minute, Z=−2.23, p=0.03, BF_10_=0.87) and no significant effect of the observers’ isolation time on the number of pins was observed (−1.20 occurrences [−3.40, 1.00] per minute, Z=−1.04, p=0.30, BF_10_=0.22 and −0.15 occurrences [−2.10, 1.80] per minute, Z=−0.169, p=0.88, BF_10_=0.16; respectively for 120 and 270 min compared to 30 min, Figure 3D and Table 10). The same effect was found when quantifying total play duration (Supp Figure 4B and Table 12).

**11.**
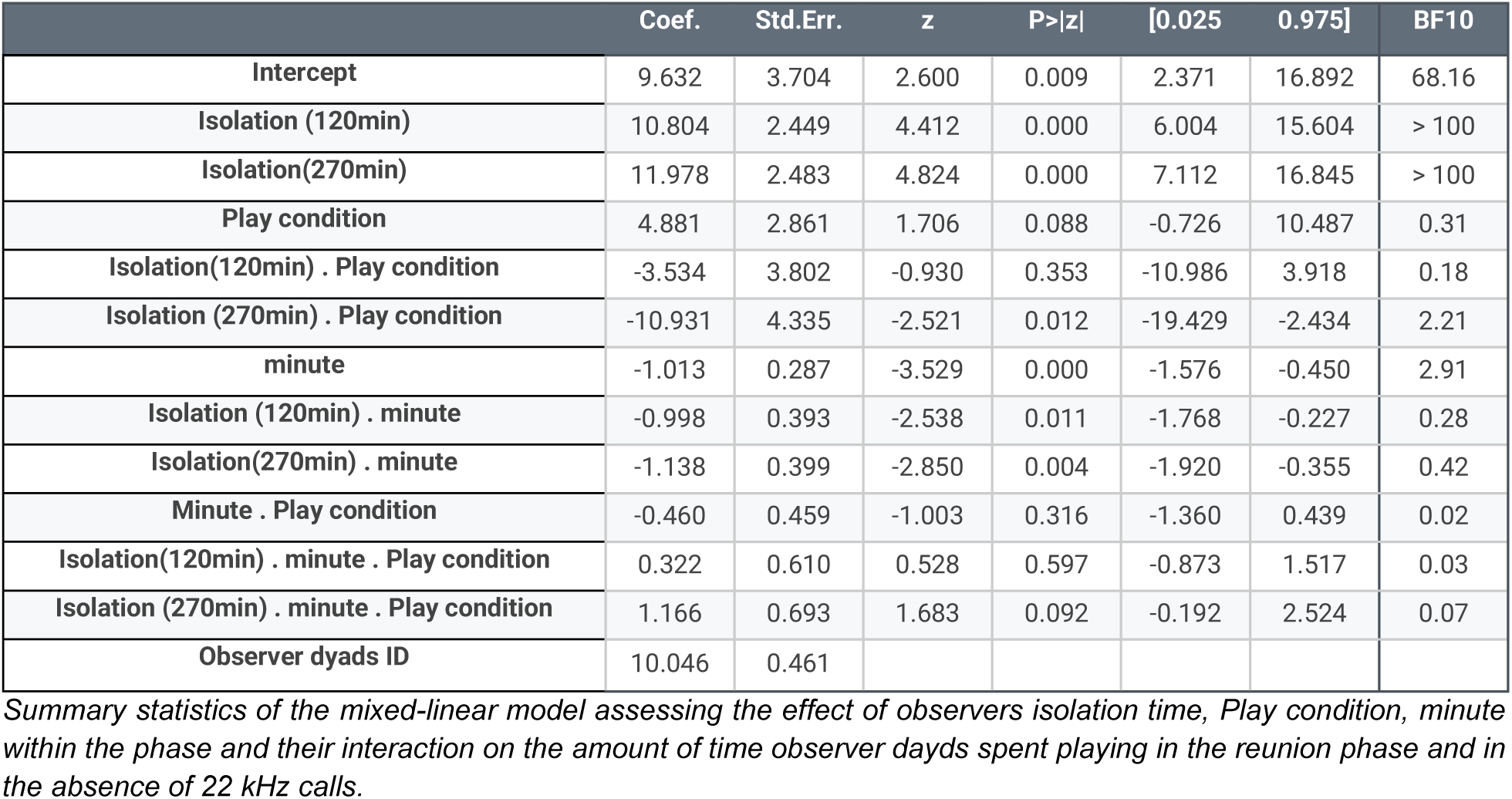
Observers Play duration - Reunion Phase - no 22KHz.

**12.**
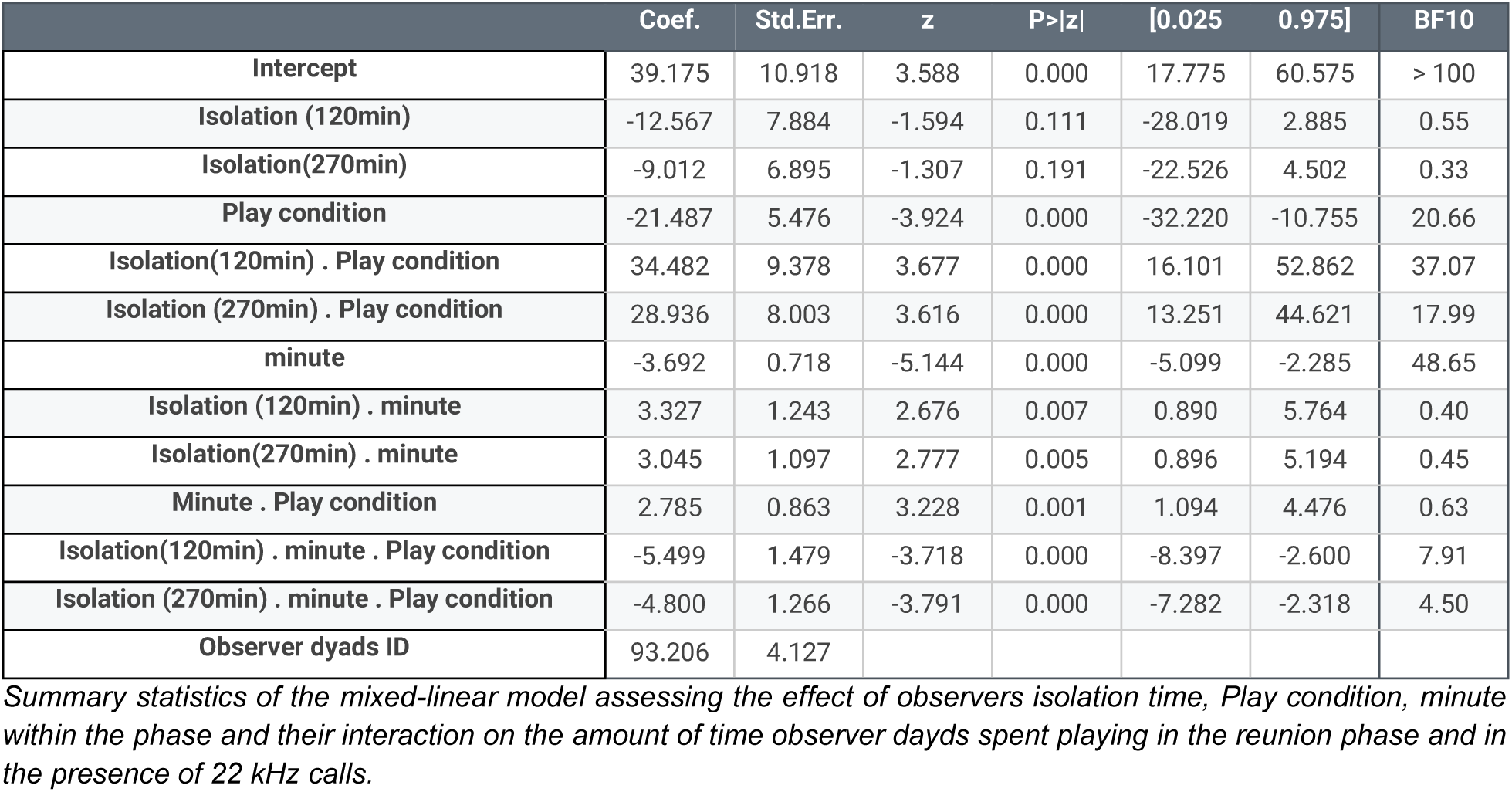
Observers play duration - Reunion Phase - 22KHz.

To assess whether the effect of Play observation on play behavior in sessions without 22 kHz calls was due to observer rats engaging in more play behavior than in sessions with 22 kHz calls, we compared the number of pins rats displayed between these sessions (Supp Figure 4 and Tables 13-14). Surprisingly, we found no effect following Play observation (+0.23 occurrences [−1.00,1.50] per minute, Z=0.37, p=0.71, BF_10_=0.28), but rather an *increased* pinning in sessions with 22 kHz calls following control observation (+3.10 occurrence [1.90,4.40] per minute, Z=4.77, p<0.001, BF_10_>100). Together with the observation of enhanced play behavior by demonstrator rats in sessions with 22 kHz calls (section 3.1 and Figure 2D), these results suggest that for sessions with 22 kHz calls emitted, Play observation did not seem to influence the behaviour of the observers.

**13.** Observers Pinning Frequency - Reunion Phase - 22KHz split Control.

13- Observers Pinning Frequency - Reunion Phase - 22KHz split Control
|  | Coef. | Std.Err. | z | P>\|z\| | [0.025 | 0.975] | BF10 |
| --- | --- | --- | --- | --- | --- | --- | --- |
| Intercept | 4.551 | 0.707 | 6.439 | 0.000 | 3.166 | 5.936 | > 100 |
| Isolation (120min) | -1.107 | 0.985 | -1.124 | 0.261 | -3.038 | 0.823 | 0.29 |
| Isolation(270min) | -0.174 | 0.866 | -0.200 | 0.841 | -1.871 | 1.524 | 0.28 |
| 22kHz calls absence | -3.140 | 0.659 | -4.767 | 0.000 | -4.432 | -1.849 | > 100 |
| Isolation(120min) . 22kHz calls absence | 2.951 | 1.072 | 2.754 | 0.006 | 0.851 | 5.051 | 3.96 |
| Isolation (270min) . 22kHz calls absence | 2.738 | 0.965 | 2.837 | 0.005 | 0.846 | 4.629 | 6.19 |
| minute | -0.502 | 0.091 | -5.525 | 0.000 | -0.679 | -0.324 | > 100 |
| Isolation (120min) . minute | 0.308 | 0.157 | 1.956 | 0.050 | -0.001 | 0.616 | 0.14 |
| Isolation(270min) . minute | 0.122 | 0.139 | 0.878 | 0.380 | -0.150 | 0.393 | 0.05 |
| Minute . Play condition | 0.332 | 0.103 | 3.228 | 0.001 | 0.131 | 0.534 | 1.22 |
| Isolation(120min) . minute . 22kHz calls absence | -0.485 | 0.171 | -2.843 | 0.004 | -0.820 | -0.151 | 0.98 |
| Isolation (270min) . minute . 22kHz calls absence | -0.382 | 0.154 | -2.474 | 0.013 | -0.684 | -0.079 | 0.45 |
| Observer dyads ID | 0.159 | 0.054 |  |  |  |  |  |

Summary statistics of the mixed-linear model assessing the effect of observers isolation time, 22kHz absence, minute within the phase and their interaction on the number of pins observer days performed playing in the reunion phase and in the control condition.

**14.** Observers Pinning Frequency - Reunion Phase - 22KHz split Play.

14- Observers Pinning Frequency - Reunion Phase - 22KHz split Play
|  | Coef. | Std.Err. | z | P>\|z\| | [0.025 | 0.975] |  |
| --- | --- | --- | --- | --- | --- | --- | --- |
| Intercept | 2.950 | 1.056 | 2.794 | 0.005 | 0.880 | 5.020 | > 100 |
| Isolation (120min) | 1.562 | 0.740 | 2.110 | 0.035 | 0.111 | 3.013 | 1.36 |
| Isolation(270min) | 2.751 | 0.579 | 4.752 | 0.000 | 1.616 | 3.885 | > 100 |
| 22kHz calls absence | -0.233 | 0.633 | -0.368 | 0.713 | -1.474 | 1.008 | 0.28 |
| Isolation(120min) . 22kHz calls absence | -0.029 | 0.952 | -0.031 | 0.975 | -1.895 | 1.836 | 0.36 |
| Isolation (270min) . 22kHz calls absence | -2.124 | 0.924 | -2.298 | 0.022 | -3.935 | -0.312 | 2.68 |
| minute | -0.273 | 0.069 | -3.938 | 0.000 | -0.408 | -0.137 | > 100 |
| Isolation (120min) . minute | -0.158 | 0.116 | -1.360 | 0.174 | -0.385 | 0.070 | 0.07 |
| Isolation(270min) . minute | -0.301 | 0.092 | -3.286 | 0.001 | -0.481 | -0.121 | 2.38 |
| Minute . 22kHz calls absence | -0.047 | 0.098 | -0.481 | 0.630 | -0.239 | 0.145 | 003 |
| Isolation(120min) . minute .22kHz calls absence | 0.006 | 0.147 | 0.042 | 0.967 | -0.282 | 0.294 | 0.05 |
| Isolation (270min) . minute . 22kHz calls absence | 0.229 | 0.143 | 1.604 | 0.109 | -0.051 | 0.509 | 0.11 |
| Observer dyads ID | 0.882 | 0.284 |  |  |  |  |  |

Summary statistics of the mixed-linear model assessing the effect of observers isolation time, 22kHz absence, minute within the phase and their interaction on the number of pins observer days performed playing in the reunion phase and in the play condition.

### 3.3 Higher overall number of 50 kHz calls emissions associated with Play condition

Finally, and despite the inability to identify the source of each call emitted, we assessed whether Play observation also influenced the overall level of 50 kHz call emissions in our paradigm. The data was also split for sessions with and without 22 kHz calls being emitted (Figure 4, Tables 19-22). Consistent with the positive association between play and 50 kHz call emission [20], we observed substantially more 50 kHz calls emitted per minute in the Play observation condition, for both the observation and reunion phase, regardless of whether 22 kHz calls were emitted (Β_no 22 kHz_: +109.63 occurrences [94.61, 124.66] per minute, Z=14.30, p<0.001, BF_10_>100, and Β_22 kHz_: +107.21 occurrences [79.32, 135.10] per minute, Z=7.53, p<0.001, BF_10_>100; Figure 4A,C). 50 kHz call emission rates decreased over time, this effect was strongest in sessions without 22 kHz call emissions(Β_no 22 kHz_: −9.03 occurrences [−11.83, −6.24] per minute, Z=−1.43, p<0.001, BF_10_>100 and Β_22 kHz_: −5.53 occurrences [−10.60, −0.47] per minute, Z=−2.14, p=0.03, BF_10_=0.30). In addition, there was little effect of the observers’ isolation time in both sessions (see Tables 19-20).

**15.** Demonstrators proximity - Obs phase No 22KHz sessions.

15- Demonstrators proximity - Obs phase No 22KHz sessions
|  | Coef. | Std.Err. | z | P>\|z\| | [0.025 | 0.975] | BF10 |
| --- | --- | --- | --- | --- | --- | --- | --- |
| Intercept | 78.130 | 6.846 | 11.412 | 0.000 | 64.711 | 91.549 | > 100 |
| Isolation (120min) | -6.458 | 4.659 | -1.386 | 0.166 | -15.589 | 2.674 | 0.47 |
| Isolation(270min) | -0.545 | 4.659 | -0.117 | 0.907 | -9.677 | 8.587 | 0.15 |
| Play condition | 5.678 | 5.386 | 1.054 | 0.292 | -4.879 | 16.235 | 0.21 |
| Isolation(120min) . Play condition | -2.485 | 7.809 | -0.318 | 0.750 | -17.790 | 12.821 | 0.20 |
| Isolation (270min) . Play condition | -1.210 | 10.610 | -0.114 | 0.909 | -22.006 | 19.586 | 0.21 |
| Observer dyads ID | 32.119 | 2.019 |  |  |  |  |  |

Summary statistics of the mixed-linear model assessing the effect of observers isolation time, play condition and their interaction on the percentage of time observers spent in the compartment closest to the demonstrators in the observation phase and in the absence of 22 kHz calls.

**16.**
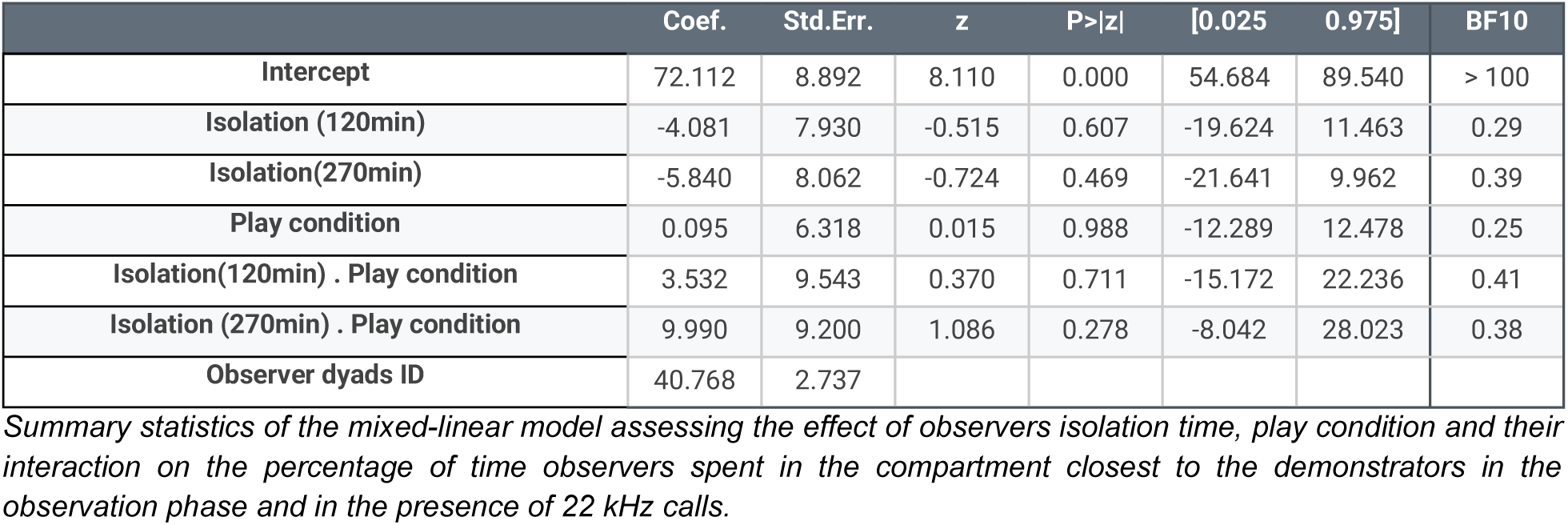
Demonstrators proximity - Obs phase 22KHz sessions.

**17.**
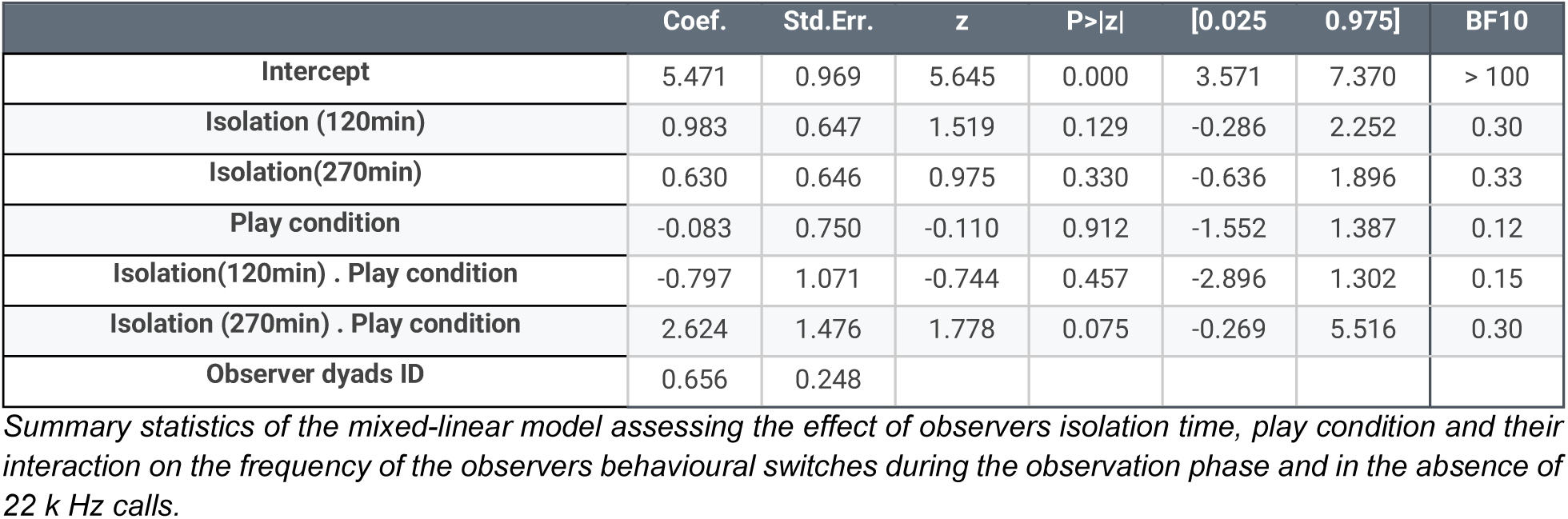
Anticipatory - Obs phase No 22KHz sessions.

17- Anticipatory - Obs phase No 22KHz sessions
|  | Coef. | Std.Err. | z | P>\|z\| | [0.025 | 0.975] | BF10 |
| --- | --- | --- | --- | --- | --- | --- | --- |
| Intercept | 5.471 | 0.969 | 5.645 | 0.000 | 3.571 | 7.370 | > 100 |
| Isolation (120min) | 0.983 | 0.647 | 1.519 | 0.129 | -0.286 | 2.252 | 0.30 |
| Isolation(270min) | 0.630 | 0.646 | 0.975 | 0.330 | -0.636 | 1.896 | 0.33 |
| Play condition | -0.083 | 0.750 | -0.110 | 0.912 | -1.552 | 1.387 | 0.12 |
| Isolation(120min) . Play condition | -0.797 | 1.071 | -0.744 | 0.457 | -2.896 | 1.302 | 0.15 |
| Isolation (270min) . Play condition | 2.624 | 1.476 | 1.778 | 0.075 | -0.269 | 5.516 | 0.30 |
| Observer dyads ID | 0.656 | 0.248 |  |  |  |  |  |

Summary statistics of the mixed-linear model assessing the effect of observers isolation time, play condition and their interaction on the frequency of the observers behavioural switches during the observation phase and in the absence of 22 k Hz calls.

**18.** Anticipatory - Obs phase 22KHz sessions.

18- Anticipatory - Obs phase 22KHz sessions
|  | Coef. | Std.Err. | z | P>\|z\| | [0.025 | 0.975] | BF10 |
| --- | --- | --- | --- | --- | --- | --- | --- |
| Intercept | 6.176 | 0.795 | 7.769 | 0.000 | 4.618 | 7.734 | > 100 |
| Isolation (120min) | -0.232 | 1.141 | -0.203 | 0.839 | -2.469 | 2.005 | 0.20 |
| Isolation(270min) | 2.150 | 1.068 | 2.012 | 0.044 | 0.056 | 4.244 | 0.43 |
| Play condition | -0.383 | 0.867 | -0.441 | 0.659 | -2.082 | 1.317 | 0.17 |
| Isolation(120min) . Play condition | -0.100 | 1.387 | -0.072 | 0.943 | -2.819 | 2.619 | 0.25 |
| Isolation (270min) . Play condition | -0.643 | 1.240 | -0.519 | 0.604 | -3.074 | 1.787 | 0.21 |
| Observer dyads ID | 0.164 |  |  |  |  |  |  |

Summary statistics of the mixed-linear model assessing the effect of observers isolation time, play condition and their interaction on the frequency of the observers behavioural switches during the observation phase and in the presence of 22 kHz calls.

**19.** 50 kHz calls - Observation phase without 22kHz sessions.

19- 50 kHz calls - Observation phase without 22kHz sessions
| Column 1 | Coef. | Std.Err. | z | P>\|z\| | [0.025 | 0.975] | BF10 |
| --- | --- | --- | --- | --- | --- | --- | --- |
| Intercept | 10.701 | 14.608 | 0.733 | 0.464 | -17.931 | 39.332 | 0.18 |
| Isolation (120min) | 13.591 | 6.565 | 2.070 | 0.038 | 0.723 | 26.459 | 0.62 |
| Isolation(270min) | 11.149 | 6.649 | 1.677 | 0.094 | -1.884 | 24.181 | 0.36 |
| Play condition | 109.632 | 7.666 | 14.302 | 0.000 | 94.608 | 124.657 | > 100 |
| Isolation(120min) . Play condition | -8.453 | 10.195 | -0.829 | 0.407 | -28.434 | 11.529 | 0.19 |
| Isolation (270min) . Play condition | 22.539 | 11.649 | 1.935 | 0.053 | -0.293 | 45.371 | 1.30 |
| minute | -1.276 | 0.892 | -1.430 | 0.153 | -3.025 | 0.473 | 0.05 |
| Isolation (120min) . minute | -1.410 | 1.222 | -1.154 | 0.248 | -3.805 | 0.985 | 0.04 |
| Isolation(270min) . minute | -1.338 | 1.241 | -1.078 | 0.281 | -3.769 | 1.094 | 0.03 |
| Minute . Play condition | -9.031 | 1.427 | -6.331 | 0.000 | -11.827 | -6.235 | > 100 |
| Isolation(120min) . minute . Play condition | 4.367 | 1.895 | 2.305 | 0.021 | 0.654 | 8.081 | 0.43 |
| Isolation (270min) . minute . Play condition | 1.257 | 2.153 | 0.584 | 0.559 | -2.963 | 5.477 | 0.04 |
| Observer dyads ID | 181.780 | 2.859 |  |  |  |  |  |

Summary statistics of the mixed-linear model assessing the effect of observers isolation time, Play condition, minute within the phase and their interaction on the number of 50 kHz calls emissions in the observation phase for sessions in the absence of 22 kHz calls.

**20.**
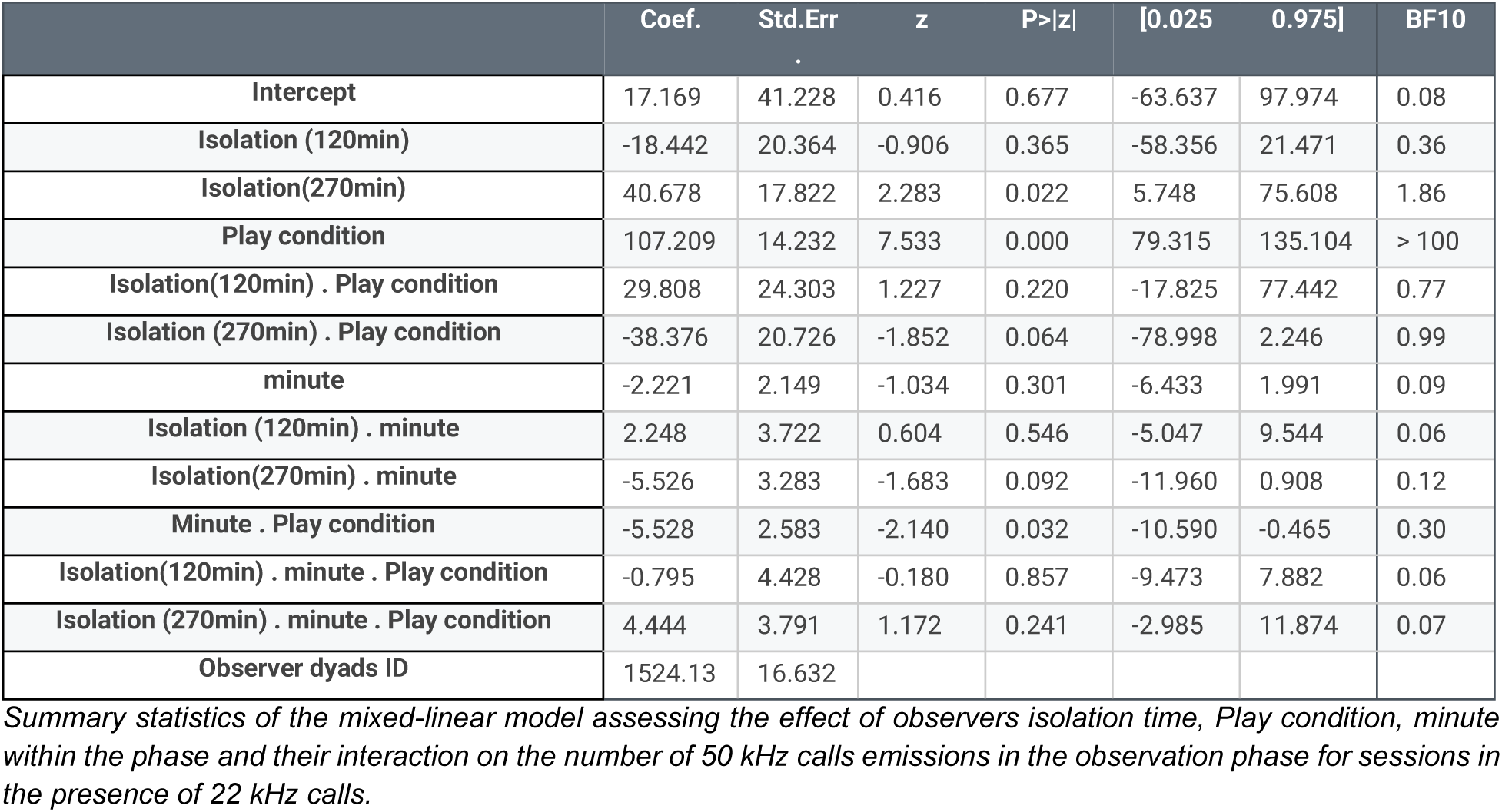
50 kHz calls - Observation phase 22KHz sessions.

20- 50 kHz calls - Observation phase 22KHz sessions
|  | Coef. | Std.Err | z | P>\|z\| | [0.025 | 0.975] | BF10 |
| --- | --- | --- | --- | --- | --- | --- | --- |
| Intercept | 17.169 | 41.228 | 0.416 | 0.677 | -63.637 | 97.974 | 0.08 |
| Isolation (120min) | -18.442 | 20.364 | -0.906 | 0.365 | -58.356 | 21.471 | 0.36 |
| Isolation(270min) | 40.678 | 17.822 | 2.283 | 0.022 | 5.748 | 75.608 | 1.86 |
| Play condition | 107.209 | 14.232 | 7.533 | 0.000 | 79.315 | 135.104 | > 100 |
| Isolation(120min) . Play condition | 29.808 | 24.303 | 1.227 | 0.220 | -17.825 | 77.442 | 0.77 |
| Isolation (270min) . Play condition | -38.376 | 20.726 | -1.852 | 0.064 | -78.998 | 2.246 | 0.99 |
| minute | -2.221 | 2.149 | -1.034 | 0.301 | -6.433 | 1.991 | 0.09 |
| Isolation (120min) . minute | 2.248 | 3.722 | 0.604 | 0.546 | -5.047 | 9.544 | 0.06 |
| Isolation(270min) . minute | -5.526 | 3.283 | -1.683 | 0.092 | -11.960 | 0.908 | 0.12 |
| Minute . Play condition | -5.528 | 2.583 | -2.140 | 0.032 | -10.590 | -0.465 | 0.30 |
| Isolation(120min) . minute . Play condition | -0.795 | 4.428 | -0.180 | 0.857 | -9.473 | 7.882 | 0.06 |
| Isolation (270min) . minute . Play condition | 4.444 | 3.791 | 1.172 | 0.241 | -2.985 | 11.874 | 0.07 |
| Observer dyads ID | 1524.13 | 16.632 |  |  |  |  |  |

Summary statistics of the mixed-linear model assessing the effect of observers isolation time, Play condition, minute within the phase and their interaction on the number of 50 kHz calls emissions in the observation phase for sessions in the presence of 22 kHz calls.

**21.**
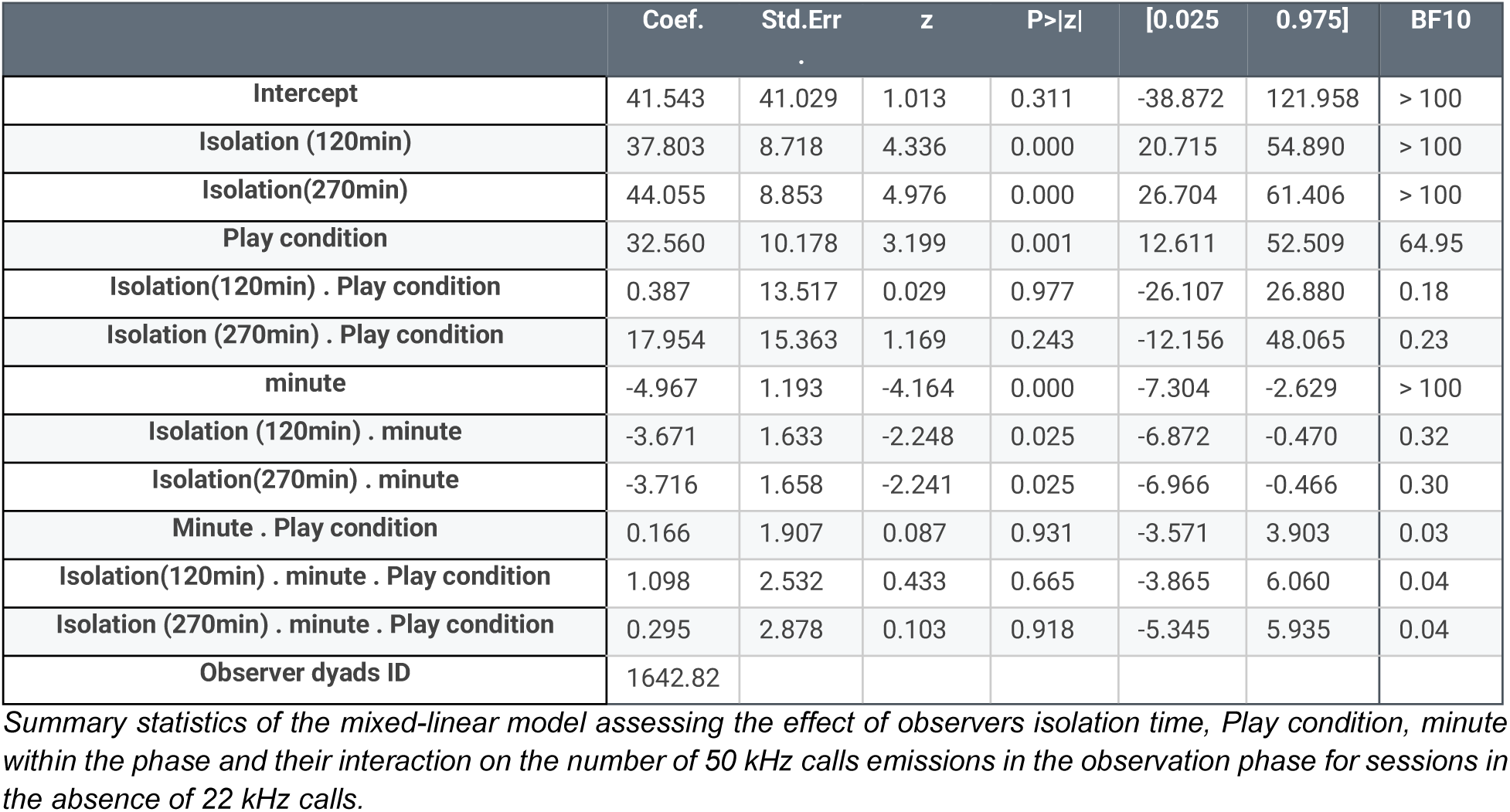
50 kHz calls - Reunion phase without 22kHz sessions.

**22.** 50 kHz calls - Reunion phase with 22kHz sessions.

22- 50 kHz calls - Reunion phase with 22kHz sessions
|  | Coef. | Std.Err | z | P>\|z\| | [0.025 | 0.975] | BF10 |
| --- | --- | --- | --- | --- | --- | --- | --- |
| Intercept | 62.944 | 22.779 | 2.763 | 0.006 | 18.298 | 107.589 | > 100 |
| Isolation (120min) | -17.835 | 21.039 | -0.848 | 0.397 | -59.071 | 23.401 | 0.30 |
| Isolation(270min) | 50.943 | 18.440 | 2.763 | 0.006 | 14.800 | 87.086 | 16.87 |
| Play condition | 25.126 | 14.710 | 1.708 | 0.088 | -3.706 | 53.958 | 0.97 |
| Isolation(120min) . Play condition | 51.089 | 25.122 | 2.034 | 0.042 | 1.849 | 100.328 | 2.65 |
| Isolation (270min) . Play condition | -1.569 | 21.417 | -0.073 | 0.942 | -43.546 | 40.407 | 0.32 |
| minute | -4.829 | 2.233 | -2.162 | 0.031 | -9.206 | -0.451 | 0.65 |
| Isolation (120min) . minute | 1.644 | 3.868 | 0.425 | 0.671 | -5.938 | 9.226 | 0.06 |
| Isolation(270min) . minute | -5.868 | 3.412 | -1.720 | 0.085 | -12.555 | 0.818 | 0.18 |
| Minute . Play condition | -1.063 | 2.684 | -0.396 | 0.692 | -6.324 | 4.198 | 0.04 |
| Isolation(120min) . minute . Play condition | -2.400 | 4.601 | -0.522 | 0.602 | -11.418 | 6.619 | 0.07 |
| Isolation (270min) . minute . Play condition | 2.004 | 3.939 | 0.509 | 0.611 | -5.717 | 9.725 | 0.06 |
| Observer dyads ID | 346.557 | 4.516 |  |  |  |  |  |

*Summary statistics of the mixed-linear model assessing the effect of observers isolation time, Play condition, minute within the phase and their interaction on the number of 50 kHz calls emissions in the observation phase for sessions in the presence of 22 kHz calls.*

During the reunion phase (Figure 4B,D, Table 21-22), we also found that 50 kHz emissions decreased with time in both sessions with and without 22 kHz calls detected (Β_no 22 kHz_: −4.97 occurrences [−7.3, −2.63] per minute, Z=−4.164, p<0.001, BF_10_>100 and Β_22 kHz_: −4.83 occurrences [−9.21, −0.45] per minute, Z=−2.16, p=0.03, BF_10_=0.65). We did find a stronger effect of the observers’ isolation time on the number of 50 kHz calls emissions (see Table 21-22), likely suggesting a stronger influence of the observers’ calls during this phase. However, mirroring the effect observed with the number of pins (section 3.2 Figure 3), we only found a statistical effect of the Play observation on the number of 50 kHz calls during the reunion phase for sessions without 22 kHz calls detected (+32.56 occurrences [12.61, 52.51] per minute, Z=3.20, p=0.001, BF_10_=63.95), despite a similar trend also being present in sessions without 22 kHz emissions (+25.13 occurrences [−3.71, 53.96] per minute, Z=1.71, p=0.088, BF_10_=0.98). Contrary to the effect of Play observation on play behaviour the effect of Play observation on 50 kHz vocalizations was independent of the observers isolation duration (see Table 21-22).

## Discussion

In this study, we aimed to capture emotional contagion induced by the observation of social play behaviour in rats. Observer rats were moderately socially deprived for 30, 120 or 270 min, watched demonstrators displaying either low or high amounts of play behavior from separate compartments, and were then reunited to freely interact after. We predicted a modulation of play behavior by the duration of social isolation of the observers, and, if emotional contagion of play occurred, heightened play following play observation compared to the control condition during the reunion phase. We used three readouts to measure the effect of play observation during the reunion phase: Total time playing, number of pins and number of 50 kHz USVs emitted. We found a convergence towards an effect akin to contagion following play observation but restricted to sessions without 22 kHz calls being emitted. Furthermore, observers showed increased pinning only when mildly socially deprived before (30 or 120 min) but not when isolated for 270 min before the sessions. We found a comparable trend for the duration of play. In addition, we detected more 50 kHz vocalizations during the reunion phase in sessions without 22 kHz calls and irrespective of the duration of observer isolation time. These results are suggestive of a subtle effect of play contagion.

### Behaviour of the demonstrator rats

In line with previous observations, demonstrator rats played more and for a longer time during the observation phase after they were socially isolated for 24 hrs than when not isolated [35,58,60]. This ensured that, during the 10 minute observation phase, observers had ample opportunity to witness play behaviour by the demonstrators in the Play observation condition, and significantly less so in the Control observation condition. In both conditions, observer animals spent more than 75% of the observation time near the demonstrator compartment, indicating that the demonstrators were a salient cue for the observers.

Of note, some demonstrator pairs could be considered better demonstrators as they played more and longer, while others displayed less social play behaviour. Indeed, some rats are more playful than others [26,49,61,62]. In paradigms of observational learning such as social transmission of food preference, social transmission of fear or social learning of foraging, the quality of the demonstrator determines subsequent performance of the observer [63–65]. This has also been demonstrated for negative emotional contagion, where observers display more freezing themselves when observing demonstrators that freeze more [66]. However, we did not find an association between the level of play of particular dyads of demonstrators and that of the observer dyads. Such a lack of a direct relationship between demonstrators and observers play could be the result of our paradigm design which strongly separated observation from the ability to express play behavior for the observers. Not being able to readily react to the play observation could have affected the experience of observer rats, thereby influencing their subsequent engagement in play. Pointing in this direction is the study of Kashtelyan and colleagues [21], where the observation of a rewarding food delivery triggered dopamine release in the ventral striatum in the observer as well as increased 50 kHz USV emission, suggesting watching a conspecific receiving a reward is initially rewarding for the observer. However, this was only true for the first trial, in subsequent trials dopamine release, 50 kHz call emission and orienting toward the reward delivery decreased to below levels of observing an empty test box. These results show that exposure to a demonstrator receiving a reward may alter the experience of the observer differently but also that multiple or extended exposures may affect the rewarding aspect of the demonstration.

### A split based on 22 kHz vocalizations

In the present study, we noticed that in some sessions 22 kHz calls were emitted. 22 kHz USVs are typically considered distress or alarm calls and are generally emitted in negatively valenced situations such as being exposed to predatory odor, and in defensive, aggressive or social conflict situations [20,67,68], although a recent study of rats in urban environments has also reported 22 kHz emissions in seemingly non-aversive situations [69]. In the current paradigm, it was impossible to pinpoint which of the rats were emitting USVs, but we could still follow their modulation over time and depending on experimental phases. 22 kHz calls emerged over the course of the 10 minute observation and reunion phases in the condition where demonstrator rats were isolated for 24 h prior to test (Play observation condition). Since 22 kHz USVs are thought to convey a negative state they could have interfered with the contagion of a positive state [18]. We therefore chose to analyse the data of sessions with and without 22 kHz emissions separately.

The 22 kHz calls could be due to one or a combination of the following: 1) an expression of a transition to a negative affective state of the observers and/or demonstrators; 2) a coordinating signal by one or both of the demonstrator or observer rats because of asymmetries in motivation to play after social isolation, for example the play interaction is considered “too rough” by one of the rats. While the first is an expression, the second is a coordinating signal to keep play rewarding [20]. Interestingly, we found that in sessions with 22 kHz calls detected demonstrator rats were highly playful. Together with the observation that the increase rate of 22 kHz calls coincide with a decline of play behaviors and 50 kHz vocalizations within the course of the 10 minute observation and reunion phases, this might point towards (1) or (2) being true; 3) During the observation phases, it may also be an expression of frustration by the observers from not being able to participate in the highly rewarding play behaviour displayed by demonstrators. In the study of Kashtelyan and colleagues [21], where observers watched demonstrator rats receive a sucrose pellet, an increase in both 50 kHz and 22 kHz USVs was found and 22 kHz USVs emerged once the demonstrators received the reward. Together with a decline in dopamine release in the ventral striatum, the authors interpreted this observation as “the observers experiencing a negative affective state akin to frustration or jealousy”. In future studies we will attempt to identify whether 22 kHz USVs are emitted by demonstrators or observers, to disentangle these interpretations using new technologies such as LiveScope [70] or a miniature microphone [71].

### Effects of play observation

In the reunion phase of sessions without 22 kHz calls, indications of a subtle contagion effect were found. Increased pinning was observed when observers were either 30 or 120 minutes but not 270 minutes socially deprived. Potentially, longer isolation times led to a high intrinsic play motivation in the observers, thus masking a potential contagion effect. This interpretation is consistent with the fact that play duration, as well as levels of pinning and of 50 kHz USV emissions increased, independently from the prior social observation with increasing observer separation time. In many studies, play deprivation is used to enhance the motivation to engage in play [24,26]. We reasoned that enhanced motivation would make the observation of play more salient and therefore would enhance the contagion effect. However, our results point to a different direction - we found stronger indications of emotional contagion with lower play deprivation lengths. In the time spent playing, we found a trend for the same subtle effect. In addition, irrespective of the length of isolation time, 50 kHz call emission was enhanced in the reunion phase after Play observation compared to the Control. These results suggest that play observation affects the internal state of the observer rat at both the behavioural and vocal levels in line with a subtle play contagion effect.

At the behavioural level, we found a stronger effect on the number of pins than on the duration of play. While play duration also encompasses play invitations (pounces) initiated by one animal, pins are an expression of mutual consent to continue the play interaction by one animal laying on its dorsal surface while the other is grooming, nuzzling or licking the ventrum [30,49]. Therefore, the fact that play observation had a stronger effect on the number of pins suggests an effect on the alignment of the motivation for the members of a dyad to engage in play.

Notably, despite the fact that the increase in 50 kHz calls emitted may come from either the observers, the demonstrators, or both, any of these could be in line with a play contagion effect, either from the play observation for the observers, or from watching playful interactions of the observers during the reunion phase for the demonstrator rats. A previous study on the contagion of a positive state induced in observer rats by witnessing tickling demonstrator rats found an increase in 50 kHz calls emitted by the observer directly after hearing the demonstrator’s 50 kHz vocalizations [16]. In line with this, identifying the emitter of specific calls would be necessary and might show a similar effect in our more naturalistic play contagion paradigm. However, 50 kHz USVs are not only considered a display of positive affect but also a means of coordinating social interactions [52,72]. Future studies are needed to disentangle their potential meaning, for example by focusing on the subtypes of 50 kHz calls that are emitted.

In our paradigm, we chose rats unfamilar to the observers as demonstrators because play behaviour is reported to be higher between strangers than cage mates [25,73]. In addition, Reinhart et al. [74] showed that when a rat in a dyadic test is satiated its level of initiating play rebounds due to the introduction of a novel, unfamiliar partner. Another study found a preference to engage in play with rats housed next to the focal rat but separated by a perforated partition preventing physical social interaction over both strangers and cage mates [75]. Moreover, it appears that the preference to play with a familiar or unfamiliar conspecific is strain dependent [76]. Thus, it would be interesting to further investigate how this balance between familiarity and novelty influences the saliency of potential social play partners in the context of play contagion.

As it is known that rats detect the positive emotional state of others through olfactory, tactile, visual, and auditory cues in rats [4,13,15,77] and that for social play behaviour specifically, auditory, tactile, olfactory and visual cues are important [38,78], another promising extension of our study would be to disentangle the multisensory information the rats exchange in our paradigm. Manipulating, in particular, visual and auditory stimuli could give us an important insight into the relevance of the different sensory modalities involved and could help to further investigate the emergence of 22 kHz vocalizations or the role of different types of 50 kHz USVs.

### Limitations of the study

To ensure that observer rats were exposed to a large number of play episodes, a 10 minute observation period was chosen. Other studies of positive emotional contagion in rats made use of food reward or tickling - mimicking social play by a human experimenter - used shorter exposure periods [16,21]. It is possible that the relatively long observation time without the possibility to engage in the behaviour could have resulted in something akin to frustration and the loss of a possible contagion effect. Another limitation of our study is that the emitting animal could not be identified, confounding the interpretation of the USV results. Through recent technological advancement it is now possible to address this factor, opening up the possibility to investigate aspects of emotional contagion on the level of an individual within a social context.

Since social play is considered a rewarding activity, we expected heightened anticipatory activity due to observing other rats engaging in social play. However, individual behaviour as well as overall behavioural switches during the observation period did not differ between Play or Control observation conditions. The same was true for proximity to the demonstrators, our measure of saliency of the reward. This could indicate that a pair of stranger rats, irrespective of whether they are engaged in play, is highly salient or that our measures of proximity and anticipatory behaviour were not sensitive enough to distinguish between the two observation conditions. In the study of Burke et al [52], anticipation of play was investigated for seven consecutive days, and anticipatory behaviour was found to be elevated at the end of this period but this was specific for adults as the juveniles did not show this effect. In our paradigm, we used three consecutive days and juvenile rats, which may explain the lack of differences found. Another output measure of anticipation is the number of 50 kHz USVs emitted [36,52]. Burke et al. [52] did find an increase in emission over days in both adult and juvenile rats. This measure of anticipation is less useful in our paradigm since demonstrator rats engaging in play also emit these vocalizations and we cannot distinguish the emitter.

A complicating factor while studying social behaviour is dyad composition. We matched pairs of demonstrators and observers based on homecage and weight but did not take partner preference of the rats into account, which is increasingly becoming recognized as an important factor for an optimal social play experience [30]. Future experiments could benefit from screening demonstrators and observers for partner preference to create more optimal dyads to capture emotional contagion.

## Conclusion

From an evolutionary perspective, the ability to recognize others’ positive emotions allows individuals to more quickly and efficiently gain access to valuable resources and safer environments by drawing on the experiences and emotions of group members, rather than relying on personal discovery [4,79,80]. Additionally, positive emotional contagion contributes to the health and well-being of individuals by reducing stress, fostering social bonding and cohesion, and enhancing the immune system [81].

In our paradigm, we found subtle indices of emotional contagion induced by the observation of demonstrator rats engaging in social play. These indices, time spent engaged in play, pins displayed and 50 kHz vocalizations emitted were dependent on several factors, such as the presence of 22 kHz calls, internal motivation of the observer and phase of experiment. Our results add to the growing body of literature investigating positive emotional contagion and provide a stepping stone towards investigating neurobiological measures of positive emotional contagion induced by social play behaviour.

## Authors contribution

Conceptualization: FML, AM, VG, CK

Data curation: FML, AM, LVK, KVR

Formal analysis: FML, AM, JP

Funding acquisition: FML, VG, CK

Investigation: FML, JP, SB

Project administration: FML

Resources: FML, VG, CK

Supervision: FML, AM, VG, CK

Visualization: FML, LVK

Writing – original draft: FML, AM, LVK

Writing – review and editing: FML, AM, LVK, JP, VG, CK

## Data

Data presented in this manuscript are available via the OpenScienceFramework, at https://osf.io/fdj62/overview?view_only=8fa33c191f244eb58c040406ccaffa6c.

## Acknowledgements

We would like to thank Nicky Markus for her help with the behavioural observations and Fleur Oogjen for her support with behavioural training..

