## Supplementary Figures for "Heightened Play following Play Observation in the Absence of 22KHz Calls in Juvenile Rats"

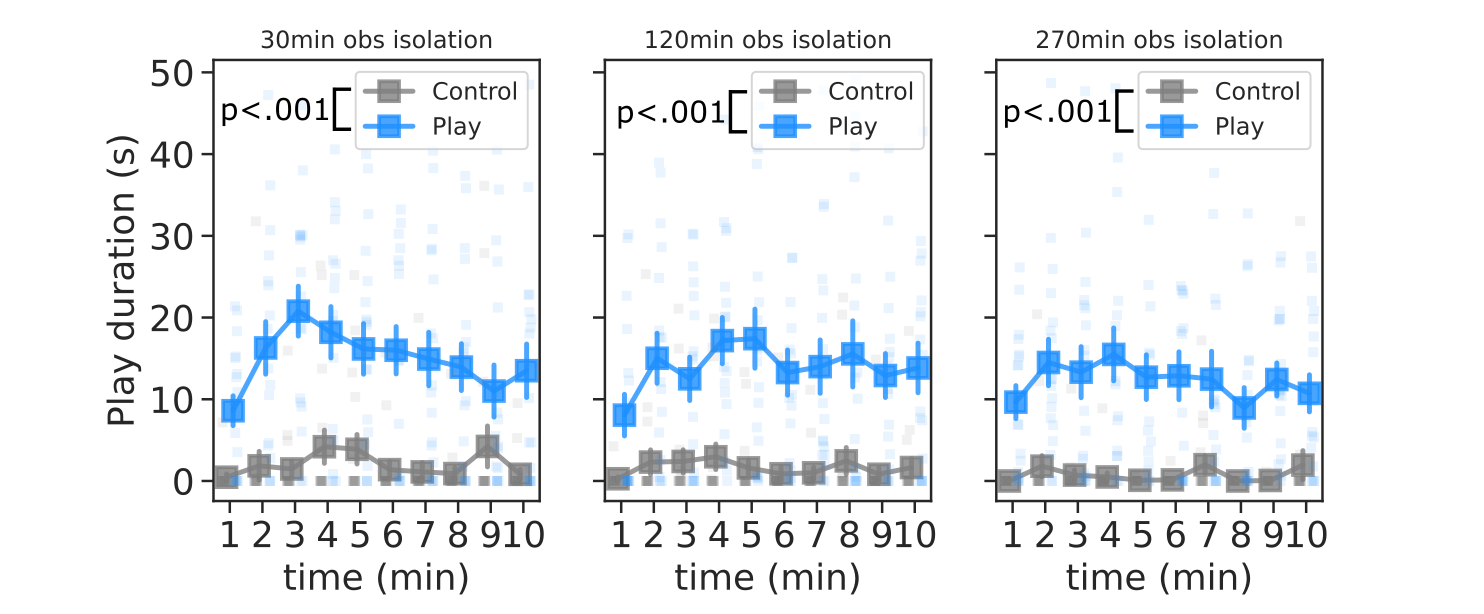

***Supplementary Figure 1: 24 h-isolated demonstrators engage in play throughout the observation phase.***

*The total amount of time spent engaging in social play displayed by the demonstrators dyads during the observation phase, separated by Control (socially housed demonstrators, grey squares) and Play (24 h-isolated demonstrators, blue squares) observation conditions and isolation time of the observers (30, 120 or 270 min prior to test). Data is shown as mean ± SEM with unconnected squares representing behavior of a single dyad.*

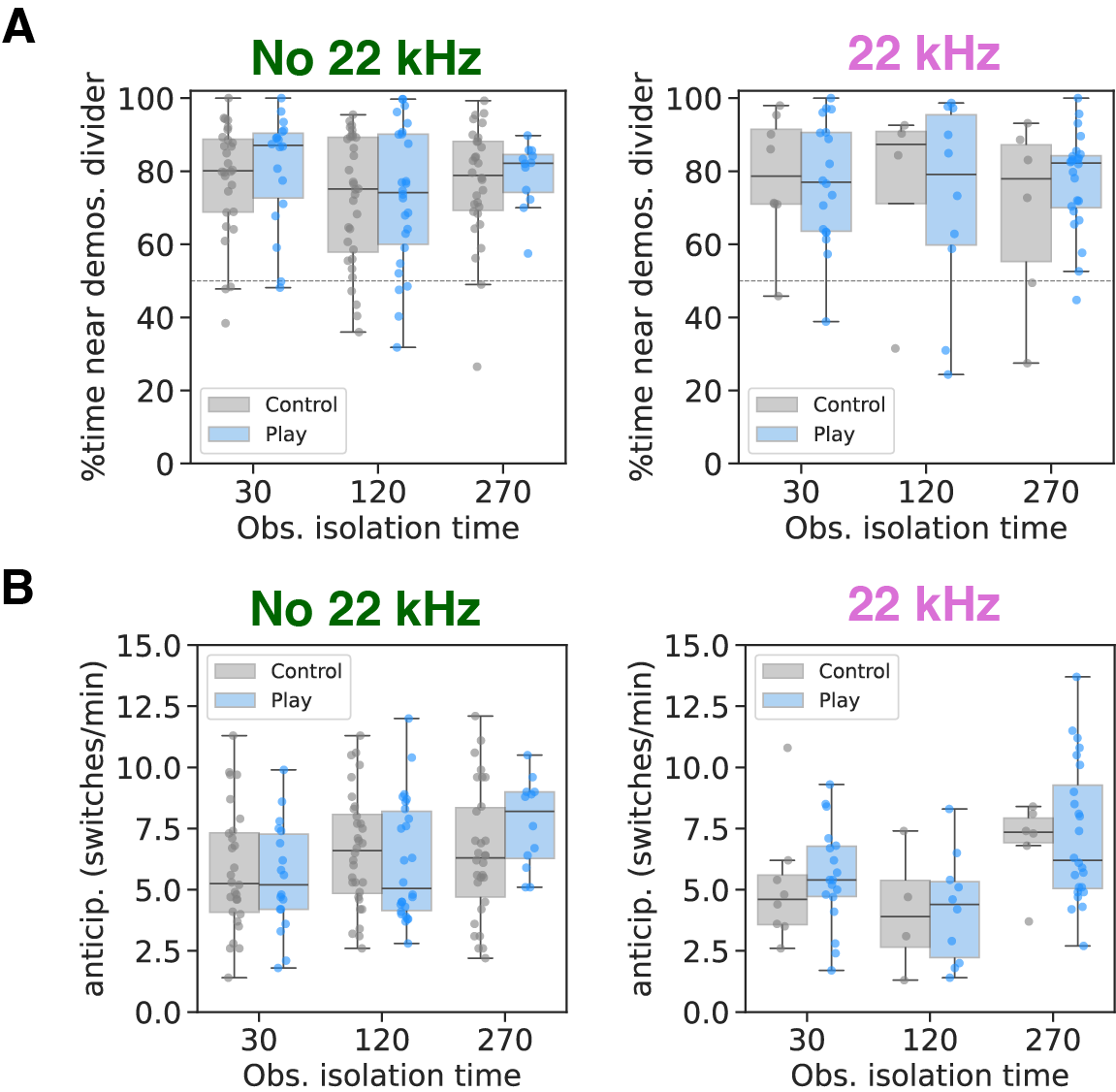

***Supplementary Figure 2: No clear effect of play observation on proximity to the demonstrator rats or anticipation of play***

***A.*** *Percentage of time spent by the observers in the half of the observer compartment closest to the demonstrators during Control (socially housed demonstrators, grey bars) and Play observation sessions (24 h-isolated demonstrators, blue bars) and per isolation time of the observers (30, 120 and 270 min prior to test) for sessions with or without 22 kHz calls being emitted. As the compartment was divided in two equally sized halves, values above 50% suggest a preference for being closer to the demonstrators.* ***B.*** *Frequency of behavioural switches per minute by the observers during Control (socially housed demonstrators, grey bars) and Play sessions (24 h-isolated demonstrators, blue bars) and per isolation time of the observers (30, 120 and 270 minutes prior to test), separated for sessions with and without 22kHz calls being emitted. Boxplots represent median and quartiles, whiskers the rest of the data distribution outside of outliers and circles indicate values for single dyads.*

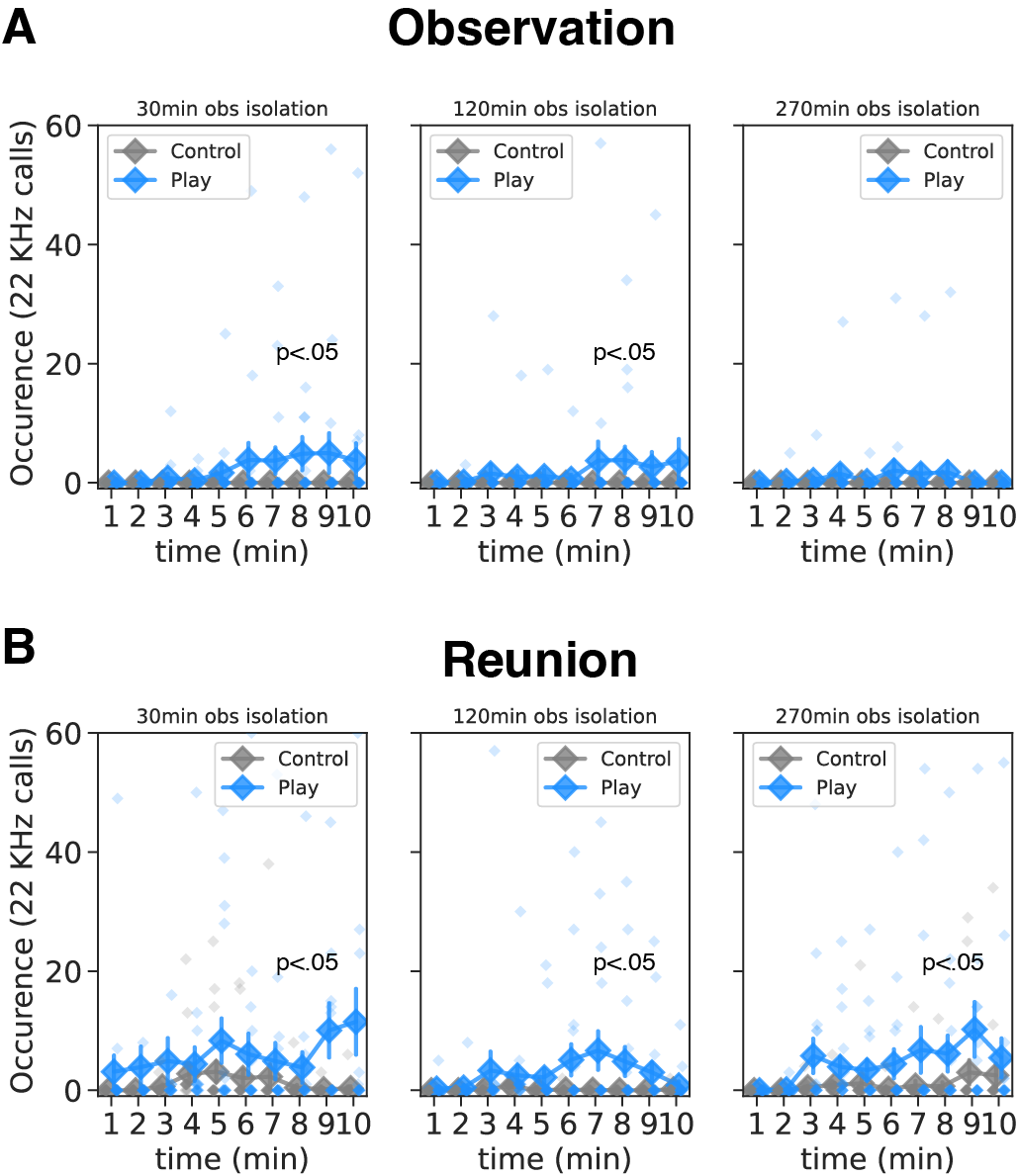

***Supplementary Figure 3: Emergence of 22 kHz call emissions in the play condition at all observers’ duration time.***

*Quantification of the number of 22 kHz calls per minute, separated by Control (socially housed demonstrators, grey diamonds) and Play (24 h-isolated demonstrators, blue diamonds) observation sessions and observers’ isolation time (30, 120, 270 min); during the observation phase (A.) and the reunion phase (B.). Connected diamonds represent the mean ± SEM, while non-connected diamonds indicate data from a single dyad.*

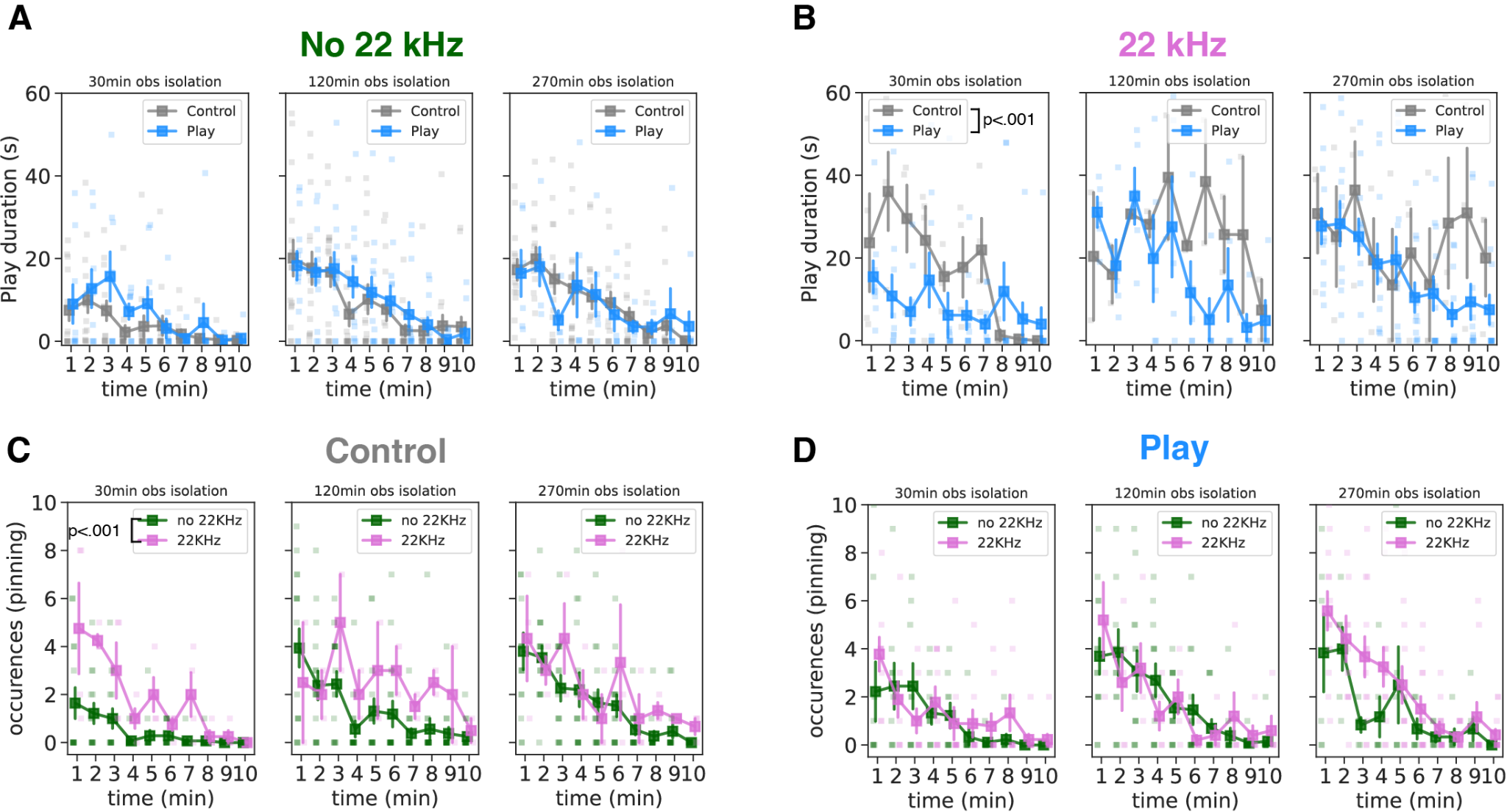

***Supplementary Figure 4: Effect of Play observation on subsequent play behavior by observer rats is masked by a heightened engagement in play behavior for sessions with 22 kHz call emission.***

***A-B.*** *Quantification of the total play duration by the observer dyads per minute in the reunion phase following observation of Control (socially housed demonstrators, grey squares) and Play (24 h-isolated demonstrators, blue squares) observation and per isolation time of the observers (30, 120 and 270 min prior to test) and separately for sessions without 22 kHz calls detected (A) and with 22 kHz calls detected (B).* ***C-D.*** *Comparison of the pins per minute displayed by observer dyads during the reunion phase between sessions with (green squares) or without 22 kHz calls detected (pink squares), separated by observers isolation time (30, 120 and 270 minutes prior to test), and following Control observation (non-socially isolated demonstrators) (C) or Play observation (24 hrs isolated demonstrators (D).*

*For all panels, connected squares represent the mean ± SEM. Non-connected squares are data from individual dyads.*
